# Mapping energy metabolism systems in the human brain

**DOI:** 10.1101/2025.03.17.643763

**Authors:** Moohebat Pourmajidian, Justine Y. Hansen, Golia Shafiei, Bratislav Misic, Alain Dagher

**Author notes:** These authors contributed equally to this work (Bratislav Misic, Alain Dagher).

## Abstract

Energy metabolism involves a network of biochemical reactions that generate ATP, utilizing substrates such as glucose and oxygen supplied via cerebral blood flow. Energy substrates are metabolized in multiple interrelated pathways that are cell- and organelle-specific. These pathways not only generate energy but are also fundamental to the production of essential biomolecules required for neuronal function and survival. How these complex biochemical processes are distributed over the cortex is integral to understanding the structure and function of the brain. Here, using curated gene sets and whole-brain transcriptomics, we generate maps of five fundamental energy metabolic pathways: glycolysis, pentose phosphate pathway, tricarboxylic acid cycle, oxidative phosphorylation and lactate metabolism. We find consistent divergence between primarily energy-producing pathways and anabolic pathways, particularly in unimodal sensory cortices. We then explore the spatial alignment of these maps with multi-scale structural and functional attributes, including metabolic uptake, neurophysiological oscillations, cell type composition, laminar organization and macro-scale connectivity. Finally, we show that metabolic pathways exhibit unique developmental trajectories from the fetal stage to adulthood. The primary energy-producing pathways peak in late childhood, while the anabolic pentose phosphate pathway shows pronounced expression in the fetal stage and declines throughout life. Together, these results highlight the rich biochemical complexity of energy metabolism organization in the brain.

## INTRODUCTION

The brain relies on substantial energy to maintain signaling and housekeeping functions. Generation of action potentials, synaptic activity, neurotransmitter release, uptake and subsequent repackaging are particularly costly processes that rely on energy production via a multitude of chemical pathways [1–3]. These fundamental and interrelated biological pathways transform nutrients to generate adenosine triphosphate (ATP), the main energy currency within the cell.

Energy metabolism is a dynamic process that changes from early development to adulthood and aging, reflecting shifts in substrate utilization and metabolic regulation to meet evolving cellular energy demands [4]. The main source of energy in the adult human brain is glucose [5, 6]. However, the brain can also utilize alternative energy resources such as lactate, ketone bodies and fatty acids under certain physiological circumstances such as intense physical activity, fasting and at specific developmental stages [4, 7, 8]. Uniquely among all organs, the brain stores virtually no energy [9]; brain cells therefore rely on constant nutrient supply from the vasculature and energy production coupled to synaptic activity. As a result, energy metabolic pathways are tightly regulated and dynamically adapt to changes in nutrient supply and demand.

Once glucose is taken up by the brain, it can be metabolized via multiple metabolic pathways, which we briefly review here (Fig. 1). Glycolysis is the first step in the breakdown of glucose. It converts one molecule of glucose to two molecules of pyruvate, while producing a net of two ATP molecules. Lactate dehydrogenases catalyze the interconversion of lactate and pyruvate, regulating the cellular redox state and enabling lactate to serve as an energy substrate that can be exported, taken up from the extracellular space, or converted to pyruvate to be used in downstream pathways [10]. Alternatively, glucose can enter the pentose phosphate pathway (PPP), an anabolic pathway essential for cellular biosynthesis. The PPP produces 5-carbon sugars for subsequent nucleotide, amino acid and neurotransmitter synthesis and generates nicotinamideadenine-dinucleotide phosphate (NADPH), an essential cofactor for lipid biosynthesis and cellular defense against reactive oxygen species (via glutathione synthesis) [7, 9, 11]. Furthermore, pentose sugars produced via PPP can be metabolized into glycolytic intermediates and subsequently converted to pyruvate by glycolytic enzymes [12, 13].

**Figure 1.**
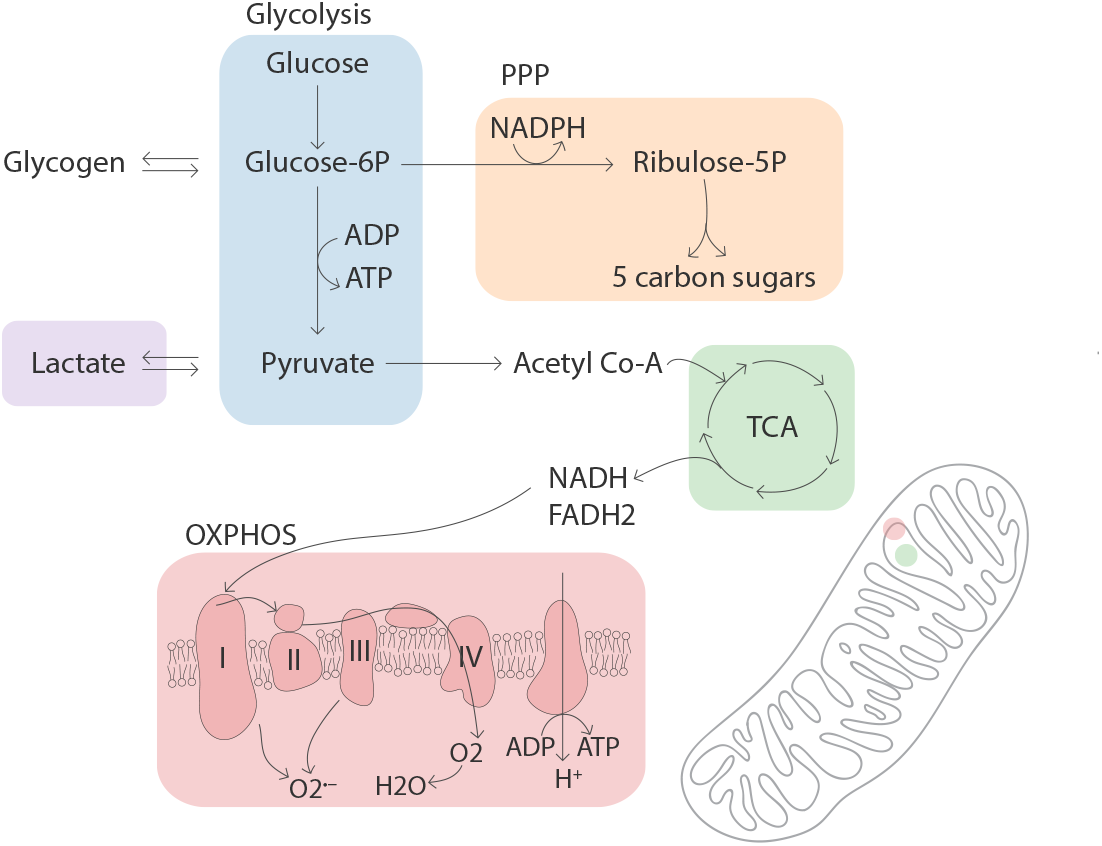
Pathways involved in brain energy metabolism. Energy metabolism refers to processes involved in energy production from nutrient molecules. Glucose is the main energy source in the brain under normal physiological conditions. Glucose entering brain cells can be utilized in three parallel pathways. It can be processed through glycolysis to produce 2 ATP and 2 pyruvate molecules (blue). Lactate dehydrogenases catalyze the interconversion of lactate and pyruvate (purple). Pyruvate is transported into mitochondria, where it enters the TCA cycle to generate high-energy electron carriers NADH and FADH2 (green), driving the complete oxidation of glucose through the mitochondrial electron transport chain (red). Glucose entering the brain can also enter the PPP. PPP is an anabolic pathway that uses glucose to produce 5-carbon sugars and NADPH, an essential co-factor used in nucleotide and lipid biosynthesis (yellow). Glucose can also be stored in the form of glycogen via glycogen synthase, a process mainly active in astrocytes. PPP, pentose phosphate pathway; TCA, tricarboxylic acid cycle; OXPHOS, oxidative phosphorylation; NADH, nicotinamide adenine dinucleotide; FADH2, Flavin adenine dinucleotide.

Pyruvate can then be shuttled to the mitochondrial matrix where it ultimately enters the tricarboxylic acid cycle (TCA) to produce high-energy electron carriers nicotinamide adenine dinucleotide (NADH) and Flavin adenine dinucleotide (FADH2). The TCA cycle is also involved in anabolic processes by providing precursors for amino acid, nucleotide and fatty acid synthesis and neurotransmitters such as glutamate and gammaaminobutyric acid (GABA) [2, 4, 14]. The high-energy electron carriers NADH and FADH2 can then enter the electron transport chain (ETC) within the mitochondrial inner membrane. The ETC is made up of four protein complexes that transfer electrons from NADH and FADH2 to molecular oxygen, producing a proton gradient across the mitochondrial membrane. ATP synthase, the fifth mitochondrial complex, uses this electrochemical gradient to produce ATP, completing the oxidative phosphorylation pathway (OXPHOS). OXPHOS enables the complete oxidation of glucose, producing *∼*15 times more ATP than glycolysis per molecule of glucose. Due to its high efficiency in ATP production, OXPHOS is regarded as the primary pathway for ATP generation in the brain [5]. Interestingly, mitochondria are also a major source of reactive oxygen species (ROS) due to electron leakage from the ETC [15, 16]. Collectively, glucose metabolic pathways are not only essential for ATP production but also play pivotal roles in anabolic processes and antioxidant homeostasis and are fundamental to cellular growth, repair, and survival.

A growing body of evidence suggests a compartmentalization of energy metabolism across the different cell types of the brain [17–19]. The differential expression of genes and enzymes, along with the selective distribution of transporters involved in energy metabolism, suggests that astrocytes favor glycolysis, whereas neurons rely more prominently on oxidative metabolism. [10, 20–23]. However, this metabolic division remains a topic of ongoing debate [6, 17] and the energy profiles of other glial cell types such as microglia and oligodendrocytes remain largely unexplored.

Multiple imaging modalities have been employed to map energy metabolism in the human brain. Positron Emission Tomography (PET) imaging has been instrumental in studying glucose uptake and oxygen consumption in the brain [24– However, PET radiotracers do not provide the biochemical resolution required to distinguish downstream glucose metabolic pathways. Magnetic resonance spectroscopy (MRS) techniques have also been used to study energy metabolism [29– MRS enables the measurement of metabolic pathway rates and metabolite concentrations, however its low sensitivity and limited spatial resolution do not allow for precise characterization and mapping of metabolic pathways in the whole brain [34, 35].

To map energy metabolism in the brain at a resolution that would allow pathway-specific insights, we employ neuroimaging transcriptomic techniques in postmortem human brains [36]. This approach provides a link between molecular data and the structural and functional architecture of the brain, allowing for a unified framework to study energy metabolism in the brain. Here, we map the distinct gene expression profile of five fundamental energy metabolism pathways across the cortex. We further explore their spatial correspondence to multi-scale structural and functional cortical features, and chart their developmental trajectories through the human lifespan.

## RESULTS

We use whole-brain microarray gene expression from the Allen Human Brain Atlas (AHBA) [38] to generate maps of five key energy metabolism pathways including: glycolysis, pentose phosphate pathway (PPP), tricarboxylic acid cycle (TCA), oxidative phosphorylation (OXPHOS) and lactate metabolism. Briefly, gene sets for each pathway were identified based on their corresponding Gene Ontology (GO) biological process [39] and Reactome pathway [40] IDs and further filtered to only retain pathway-annotated genes included in both databases. Pathway IDs and final gene sets used to produce the energy maps are provided in Table S1. Microarray transcriptome data were retrieved using the *abagen* package and parcellated into the Schaefer-400 cortical atlas ([37, 41], https://abagen.readthedocs.io/). Expression was then averaged across all genes to produce a mean gene expression map for each energy pathway.

### Mapping metabolic pathways using gene expression

Fig. 2a (left panel) shows a Venn diagram of the final number of genes included in each energy pathway expression matrix and their overlap. Note that the final gene sets for each pathway contain fewer genes compared to the original gene sets retrieved from the GO and Reactome databases, as some of the genes are not present in the AHBA or did not meet quality control criteria (see *Methods*). Importantly, there is minimal overlap between the different energy pathway gene sets, which allows the reconstruction of distinct maps that represent each pathway individually, facilitating their study in relative isolation despite their inherent interconnectivity. To assess the correspondence between metabolic pathway maps, we first compute spatial correlations among them. Fig. 2a (middle panel) shows the correlated gene expression for genes in all energy pathways across 400 cortical regions, and Fig. 2a (right panel) shows global spatial correlations among mean energy maps.

**Figure 2.**
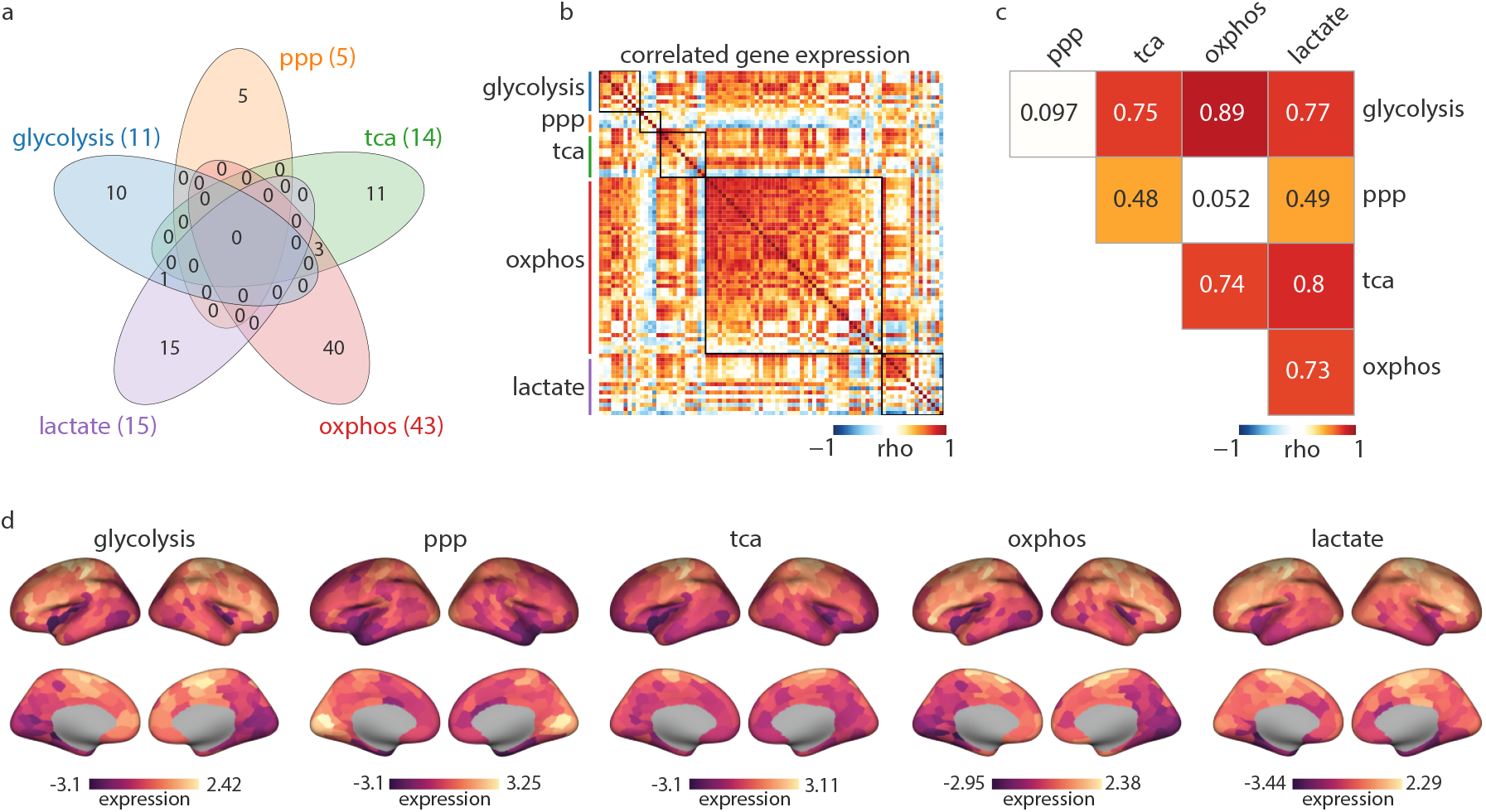
Brain maps of energy metabolism pathways. (a) Left: the Venn diagram shows the final number of genes in each pathway gene expression matrix and their overlap. Gene sets have minimal overlap; there are three genes shared between the TCA and OXPHOS belonging to the succinate dehydrogenase complex that is active in both pathways. *PFKFB2*, which encodes a regulatory enzyme of the glycolytic pathway, is shared between the glycolysis and lactate pathways. (b) Heatmap depicts the pairwise correlation of all genes included in the energy pathways across 400 regions in the Schaefer parcellation [37]. (c) Spearman’s correlation among mean expression energy maps. (d) Energy pathway maps. Color bar shows z-scored mean expression values across all genes in each pathway. For the subcortical energy pathway profiles see Fig. S3. ppp: pentose phosphate pathway; tca: tricarboxylic acid cycle; oxphos: oxidative phosphorylation; lactate: lactate metabolism and transport.

Glycolysis and OXPHOS maps show the strongest correlation (rho = 0.89, *p*_spin_ = 1 *×* 10^*−*4^) consistent with the fact that they are part of an integrated sequence of events in the oxidation of glucose [42, 43]. The lowest correlations are observed between the PPP map and the glycolysis and OXPHOS maps (rho = 0.097, *p*_spin_ = 0.79; rho = 0.052, *p*_spin_ = 0.91, respectively), in line with the role of PPP as a primarily biosynthetic pathway rather than one directly implicated in energy production [13]. Conversely, given the PPP’s function in cellular defense against ROS, it could be anticipated that regions with high oxidative metabolism would exhibit relatively elevated PPP activity. However, the PPP gene set predominantly includes genes involved in its anabolic nonoxidative branch (i.e. *PRPS2, RBKS, RPEL1, RPIA*), responsible for the production of 5-carbon sugars [6, 44]. Indeed, when we look into the *PGD* gene, the only gene representing the oxidative branch in our PPP gene set which also directly catalyzes the NADPH-producing reaction, we find significant spatial correlations with glycolysis and OXPHOS maps (glycolysis: rho = 0.75, *p*_spin_ = 1 × 10^*−*4^; OXPHOS: rho = 0.71, *p*_spin_ = 1 × 10^*−*4^; Fig. S1).

The PPP map shows a moderate correlation with the TCA map (rho = 0.48, *p*_spin_ = 0.007), potentially reflecting their shared roles in supporting cellular anabolic processes, and highlighting cortical regions with greater biosynthesis demands. Furthermore, the TCA map also shows a strong alignment with the glycolysis and OXPHOS maps (rho = 0.75, *p*_spin_ = 0.005; rho = 0.74, *p*_spin_ = 0.004, respectively). This highlights the TCA’s dual role in both energy production and cellular biosynthesis [14, 19, 45]. The lactate map shows strong correlations with glycolysis (rho = 0.77, *p*_spin_ = 1 *×* 10^*−*4^), TCA (rho = 0.8, *p*_spin_ = 2 *×* 10^*−*4^), and OXPHOS (rho = 0.73, *p*_spin_ = 1 × 10^*−*4^), likely reflecting lactate’s role as a versatile intermediate in brain energy metabolism. Lactate, produced either via aerobic glycolysis or taken up from the vasculature, is converted to pyruvate by lactate dehydrogenase and readily utilized in the TCA cycle [10, 42]. The spatial alignment observed between lactate and the other energy pathway maps underscores its role in shuttling metabolic substrates across pathways, linking glycolysis and oxidative metabolism. Collectively, these results highlight the dependencies and interplay between energy pathways. In the following subsection we investigate the regional heterogeneity of these pathways and their enrichment across structural and functional systems.

### Regional heterogeneity of metabolic pathways

How are metabolic pathways distributed across the cortex? Fig. 2b shows that energy pathway gene expression is regionally heterogeneous. Glycolysis and OXPHOS exhibit greater expression in motor and prefrontal cortex, and lower expression in visual and parietal association cortex. In contrast, the PPP map shows greater expression in the visual cortex. The TCA map shows a particularly higher expression in somato-motor regions (Fig. 2b, Fig. S2). Across the subcortical regions, energy maps consistently show greater expression in the thalamus and lower expression in the amygdala (Fig. S3).

To investigate how the spatial patterning of metabolic pathways aligns with the structural and functional organization of the brain, we estimate their enrichment in four atlases: (1) cytoarchitecture (von Economo-Koskinas classes); [46, 49]), (2) synaptic and laminar architecture (Mesulam classes; [47, 50, 51]), (3) unimodal-transmodal hierarchy [48, 50], and (4) intrinsic functional networks (Yeo-Krienen networks; [37, 52]). For each network class, the average expression of parcels falling into that class was calculated and tested against a distribution of 10 000 spatial autocorreltaionpreserving null. Across the seven von Economo-Koskinas cytoarchitectonic classes, all energy pathway maps except for the PPP have significantly higher expression in the primary motor cortices (glycolysis: *p*_spin_ = 0.01; TCA: *p*_spin_ = 8 *×* 10^*−*4^; OXPHOS: *p*_spin_ = 0.008; lactate: *p*_spin_ = 0.002; Fig. 3). The PPP map shows significantly greater average expression in the primary sensory cortex (*p*_spin_ = 1 *×* 10^*−*4^). The insular cortex has the lowest expression across all energy maps with PPP, TCA and lactate maps showing significantly lower values (PPP: *p*_spin_ = 0.003; TCA: *p*_spin_ = 4 *×* 10^*−*4^; lactate: *p*_spin_ = 0.04).

**Figure 3.**
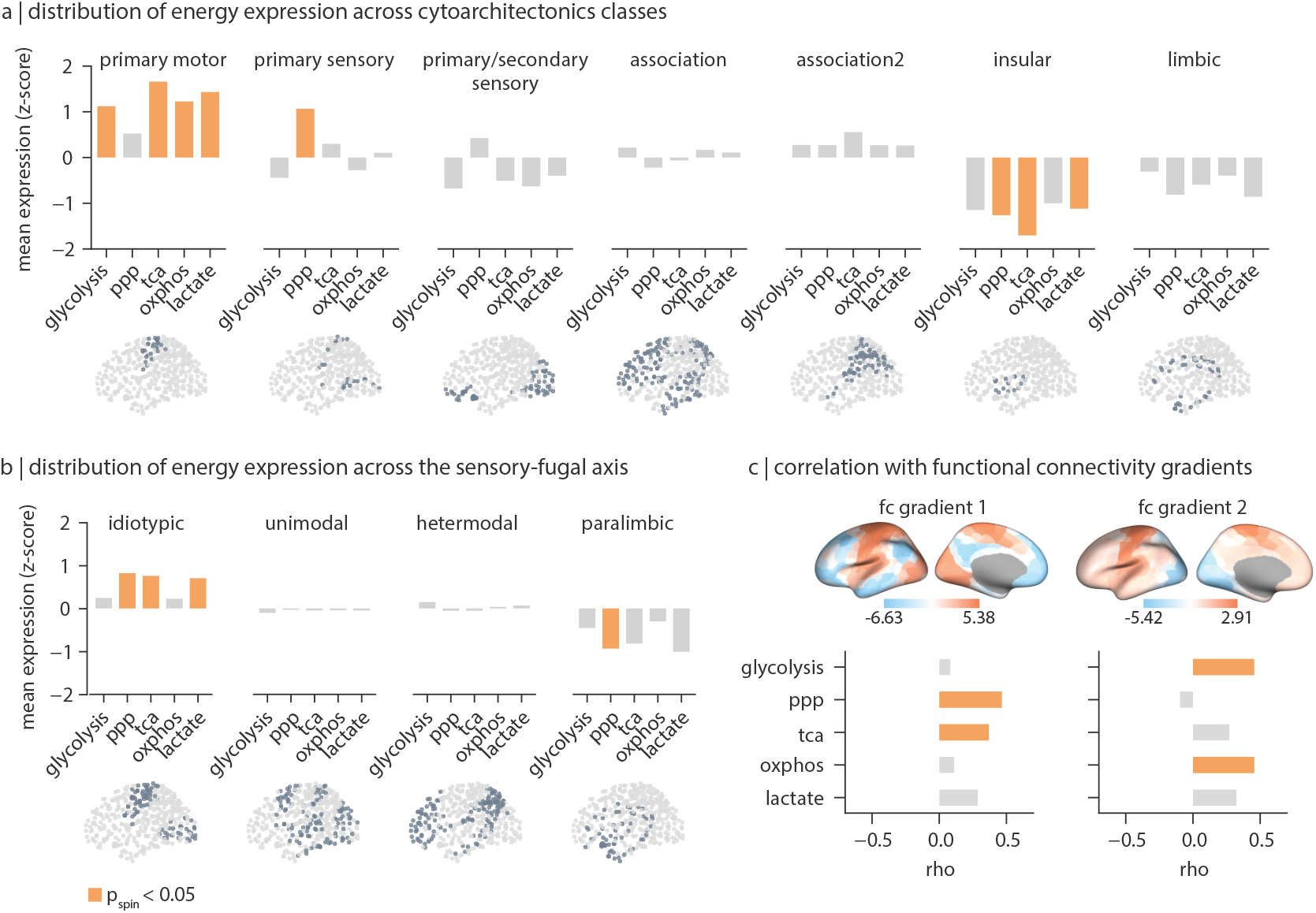
Distribution of energy pathway maps across structural and functional networks in the cortex. For each energy map, average expression values for parcels falling into each structural and functional class was calculated. (a) Distribution of energy pathway mean expression across seven von Economo cytoarchitectonic classes [46]. (b) Distribution of energy maps across the sensory-fugal axis of information processing [47]. Bars represents observed average expression for each energy map in each network class. (c) Spearman’s correlation between energy maps and the first two functional connectivity (FC) gradients in the human cortex [48]. Highlighted bars indicate statistical significance when tested against 10 000 spatial-autocorrelation preserving nulls (*p*_spin_ < 0.05). Brain plots visualize regions included in each network class. ppp, pentose phosphate pathway; tca, tricarboxylic acid cycle; oxphos, oxidative phosphorylation; lactate: lactate metabolism and transport; fc, functional connectivity.

To investigate how the energy maps align with the broader cortical hierarchy and functional organization, we further estimated their enrichment across the Mesulam sensory-fugal hierarchy and the first two principal gradients of resting state functional connectivity (FC) [48]. Along the Mesulam sensory-fugal axis, the PPP, TCA and lactate maps exhibit a similar enrichment, with significantly greater expression in idiotypic areas (PPP: *p*_spin_ = 0.002; TCA: *p*_spin_ = 0.002; lactate: *p*_spin_ = 0.001), and overall lower expression in the paralimbic regions. Glycolysis and OXPHOS maps do not show significant variations across this hierarchy. Likewise, the first FC gradient (FC1) significantly correlates with the PPP and TCA maps (PPP: rho = 0.47, *p*_spin_ = 5 *×* 10^*−*4^; TCA: rho = 0.37, *p*_spin_ = 0.03) corresponding to overall greater expression in primary cortices and lower expression in higher order association areas. The second FC gradient (FC2), which differentiates within the primary cortices, correlates significantly with the glycolysis (rho = 0.47, *p*_spin_ = 0.016) and OXPHOS maps (rho = 0.46, *p*_spin_ = 0.01), reflecting greater expression in motor regions and lower expression in the occipital cortex. Fig. S4 shows similar findings using the finer delineation of these functional patterns according to the Yeo-Krienen intrinsic networks. Collectively, these results highlight the heterogeneous distribution of energy metabolism pathways across the structural and functional organization of the cortex, and point to a consistent dichotomy between pathways primarily involved in ATP production (glycolysis and OXPHOS) and the anabolic PPP. The TCA cycle, which contributes to both oxidative metabolism and anabolic processes, integrates features of both metabolic functions.

### Energy gradients in the visual cortex

The visual cortex stands out in our analyses as one of the regions in which energy pathways are most differentiated, with greater expression of PPP and lower expression of glycolysis and OXPHOS. However, previous research characterizes the visual cortex as having higher glucose and oxygen consumption [54, 55]. This discrepancy may be due to the underlying heterogeneity within the visual cortex, where individual sub-regions exhibit distinct metabolic profiles that are not captured when analyzed as a single structure. Here, we investigate the distribution of the energy maps across the information processing hierarchy of the visual cortex as defined by the Glasser atlas ([53], Table S2). TCA, OXPHOS and lactate show greater expression in the dorsal stream and lower expression in the ventral stream (Fig. 4). Glycolysis also exhibits greater expression in the dorsal stream but lower expression in the primary visual cortex (V1). This is potentially in line with previous reports of higher cytochrome oxidase reactivity in the magnocellular layers of the lateral geniculate nucleus relative to the parvocellular layers [56]. The dorsal stream is thought to be primarily driven by the magnocellular pathway, which processes high temporal frequencies, while the ventral pathway is dominated by the parvocellular pathway that responds to low temporal frequency [56– The differences in energy pathway expression may reflect the unique energy demands for visual attributes processed via the dorsal versus ventral streams. The PPP pathway, on the other hand, shows higher expression in the primary visual cortex. Together, these findings suggest that distinct components of the visual information processing hierarchy impose unique demands on energy metabolism.

**Figure 4.**
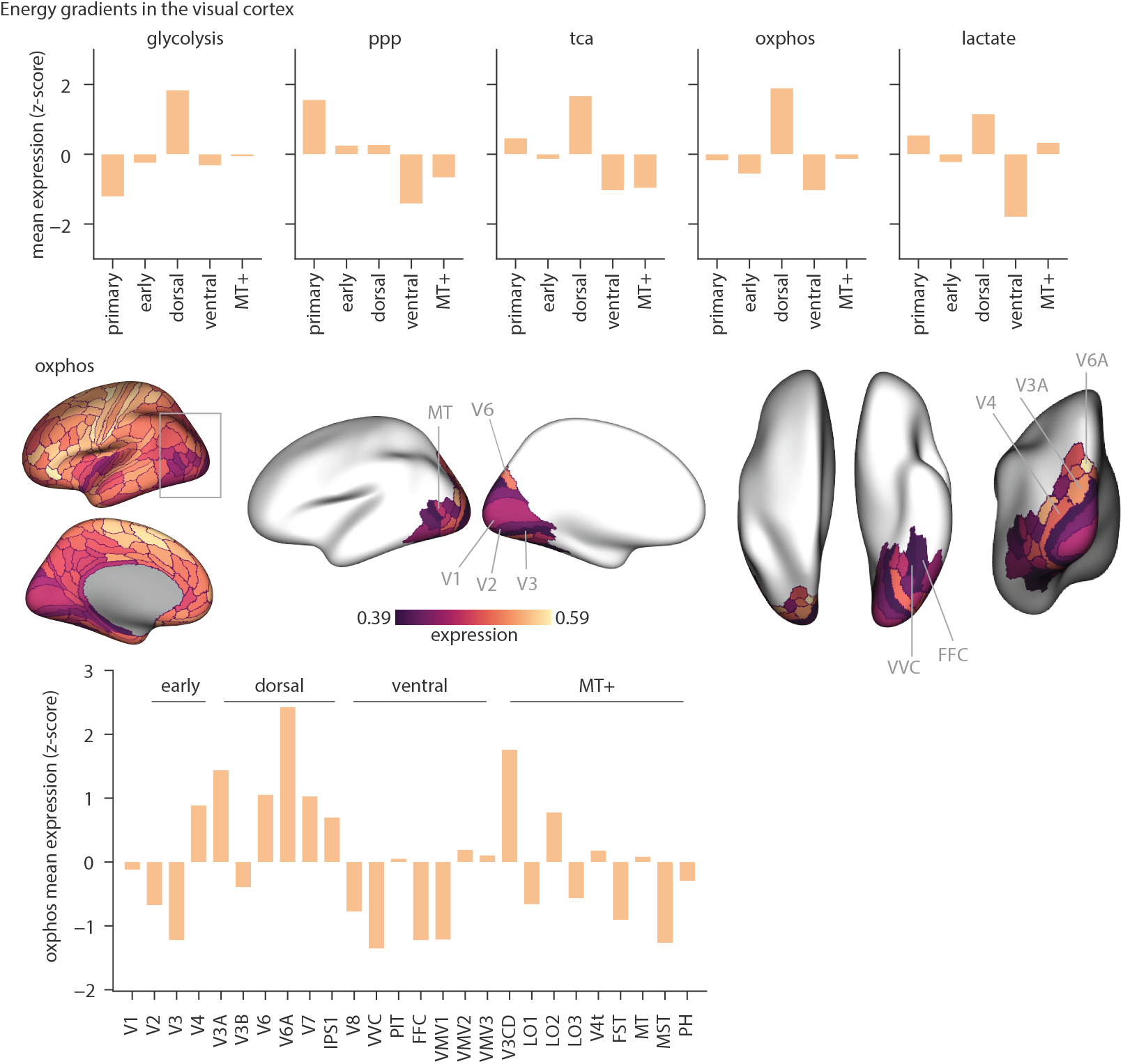
Spatial distribution of energy pathways within the visual cortex. Energy maps were produced using the Glasser parcellation and the visual parcels were grouped according to information processing hierarchy detailed in [53] (see Table S2). Top: enrichment of energy maps across the visual hierarchy. Bars represents z-scored mean gene expression of each pathway, averaged across all ROIs in each hierarchy level. Middle: distribution of OXPHOS mean gene expression across the visual cortex shown for the left hemisphere. From left to right: lateral, medial, dorsal, ventral and posterior views. Color bar represents the range of expression values within the visual cortex. Bottom: distribution of oxphos expression across individual visual ROIs. Bars represent z-scored mean expression of OXPHOS genes. dorsal, dorsal stream; ventral, ventral stream, MT+, MT+ complex and neighboring visual areas; ppp, pentose phosphate pathway; tca, tricarboxylic acid cycle; oxphos, oxidative phosphorylation; lactate: lactate metabolism.

### Energy correlates of multi-scale cortical features

The heterogeneity of energy pathway profiles likely arises from the distinct molecular and cellular properties of different cortical regions at the microscale, as well as neurophysiological and network-level attributes at the macroscale. Cortical regions exhibit variability in their glucose uptake and oxygen consumption, as shown by molecular imaging. Neurophysiological activity associated with signaling (i.e., action potentials and and synaptic transmission) are energy-intensive and account for the majority of cortical energy demand [1, 5, 8, 60]. Cellular specialization further shapes energy metabolism, as glial and neuronal populations exhibit distinct metabolic profiles. Here, we characterize the spatial alignment of the energy pathways with a collection of maps corresponding to: (1) molecular imaging of metabolic uptake from PET [54], (2) neurophysiological oscillations from magnetoencephalography (MEG) [61], (3) cell type composition, (4) laminar organization [62, 63] and (5) connectivity metrics (Fig. 5).

**Figure 5.**
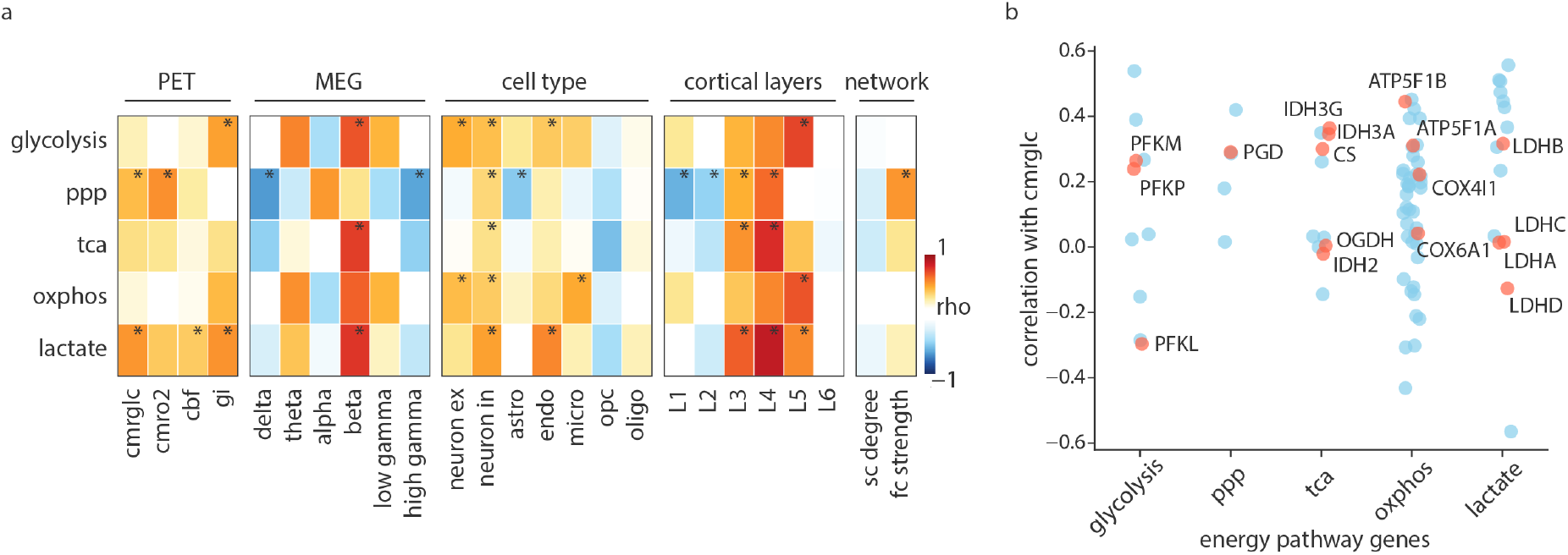
Spatial correspondence of energy maps with multi-scale cortical annotations. (a) Spearman’s correlation between energy maps and a set of brain maps including *in vivo* metabolic PET imaging [54], neurophysiological oscillations [61], cell type composition, cortical laminar thickness [62] and network connectivity attributes [67]. Colors depict the strength of the correlation between pairs of brain maps. Asterisks indicate statistical significance when tested against a distribution of 10 000 spatial-autocorrelation preserving nulls after FDR-correction for multiple comparisons. (b) Correlation between individual genes in the energy pathways and PET-derived glucose uptake. Spearman’s correlation was calculated between gene expression values and the CMR_*glc*_ map. Red dots indicate genes encoding rate-limiting and key catalytic enzymes. CMR_*glc*_, cerebral metabolic rate of glucose; CMR__*O*2__, cerebral metabolic rate of oxygen; gi, glycolytic index; cbf, cerebral blood flow. Cell types: astro, astrocyte; neuron ex, excitatory neuron; neuron in, inhibitory neuron; endo, endothelial cell; micro, microglia; opc, oligodendrocyte precursor cells; oligo, oligodendrocyte; sc, structural connectivity; fc: functional connectivity.

The CMR_*glc*_ map, which corresponds to glucose uptake, does not significantly correlate with glycolysis (rho = 0.22, *p*_*spin*,*FDR*_ = 0.63) and OXPHOS maps (rho = 0.16, *p*_*spin*,*FDR*_ = 0.76), however, it shows significant correlations with the lactate (rho = 0.53, *p*_*spin*,*FDR*_ = 0.005) and PPP (rho = 0.42, *p*_*spin*,*FDR*_ = 0.02) maps. Given that the PPP is generally characterized by minimal flux relative to other energy-producing pathways [6, 12, 44], this strong correlation with CMR_*glc*_ is unexpected. However, it has been shown that glucose flux through the PPP is largely underestimated [64, 65]. Also, contrary to what we expected, the CMR_*O*2_ map, representing oxygen consumption in the brain, does not correlate with the OXPHOS map (rho = *−*0.01, *p*_*spin*,*FDR*_ = 0.95). There is a moderate alignment between CMR_*O*2_ and the PPP map (rho = 0.54, *p*_*spin*,*FDR*_ = 0.002), potentially highlighting elevated cellular repair processes in regions with greater oxygen consumption and consequently higher oxidative damage. The glycolytic index (GI) map shows significant positive correlations with the glycolysis (rho = 0.49, *p*_*spin*,*FDR*_ = 0.047) and lactate (rho = 0.53, *p*_*spin*,*FDR*_ = 0.04) maps, in line with GI being a measure of aerobic glycolysis in the brain [54, 66]. Unexpectedly, the PPP map shows no correlation with the GI map, despite contributing to non-oxidative glucose consumption. However, this may suggest that the PPP is not a major determinant of GI and that lactate production primarily drives aerobic glycolysis. A multitude of steps bridge gene transcription and metabolic activity. This is evident when examining the gene-wise spatial alignment with CMR_*glc*_, revealing a wide range of correlations within each energy pathway (Fig. 5b). Collectively, these results highlight the complexity of downstream metabolic pathways activated after glucose uptake, perhaps capturing a different aspect of the underlying biology.

To examine how oscillatory activity is supported by energy metabolism pathways, we looked at the correspondence between energy maps and the six canonical MEG power bands. The energy maps primarily correlate with the beta band, reflecting their greater expression in the motor cortex (glycolysis: rho = 0.69, *p*_*spin*,*FDR*_ = 0.03; OXPHOS: rho = 0.65, *p*_*spin*,*FDR*_ = 0.07; TCA: rho = 0.74, *p*_*spin*,*FDR*_ = 0.04; lactate: rho = 0.77, *p*_*spin*,*FDR*_ = 0.005). We next explored the correspondence between energy maps and cell type composition. Excitatory neurons show moderate correlations with the glycolysis (rho = 0.46, *p*_*spin*,*FDR*_ = 0.001), and OXPHOS (rho = 0.42, *p*_*spin*,*FDR*_ = 0.006) maps, in line with the higher energy demand of these principal cortical neurons [3, 5, 68]. Inhibitory neurons show significant correlations across all energy maps (glycolysis: rho = 0.42, *p*_*spin*,*FDR*_ = 0.001; OXPHOS: rho = 0.37, *p*_*spin*,*FDR*_ = 0.001; TCA: rho = 0.3, *p*_*spin*,*FDR*_ = 0.003; Lactate: rho = 0.5, *p*_*spin*,*FDR*_ = 0.001; PPP: rho = 0.33, *p*_*spin*,*FDR*_ = 0.002). This can be attributed to both the greater energy demand and need for cellular repair processes due to their fast-spiking activity (see *Discussion*). Endothelial cells show significant positive correlations with the glycolysis (rho = 0.38, *p*_*spin*,*FDR*_ = 0.014) and lactate maps (rho = 0.56, *p*_*spin*,*FDR*_ = 0.001), in line with the proposed glycolytic nature of these cells [69, 70]. Microglia significantly correlate with the OXPHOS map (rho = 0.47, *p*_*spin*,*FDR*_ = 0.003), consistent with their reliance on oxidative metabolism [2, 71, 72]. The PPP map however, exhibits a negative correlation with the astrocyte map and no significant correlation with microglia, OPC and oligodendrocytes, contrary to the higher activity of this pathway in glial cells [12, 73, 74].

Given the diverse cellular composition and the distinct circuitry of cortical layers, we next sought to investigate the energy metabolism profile of cortical laminar organization. The PPP, TCA and lactate maps show the greatest alignment with cortical layer 4 (PPP: rho = 0.64, *p*_*spin*,*FDR*_ = 0.001), TCA: rho = 0.79, *p*_*spin*,*FDR*_ = 0.001, lactate: rho = 0.87, *p*_*spin*,*FDR*_ = 0.001), hinting at a greater need for fast energy supply and cellular biosynthesis in this sensory input layer. The glycolysis and the OXPHOS maps show significant alignment with the cortical layer 5 (glycolysis: rho = 0.73, *p*_*spin*,*FDR*_ = 0.001, OXPHOS: rho = 0.69, *p*_*spin*,*FDR*_ = 0.001). This could hint to the greater metabolic demand of large pyramidal cells with their extensive subcortical projections [75, 76] Overall, energy maps exhibit diverse alignments across the different cell type and the cortical laminar organization, potentially pointing to the underlying compartmentalization of energy metabolism.

The relationship between energy metabolism and network connectivity can offer insight into the metabolic demands of topological hubs within cortical networks. Here, we find that the FC strength correlates positively with the PPP map (rho = 0.52, *p*_spin_ = 0.015), suggesting greater anabolic demand in the functional hubs and in line with the higher glycolytic index and plasticity of these hub regions [66, 77]. However, the FC strength does not correlate with glycolysis nor OXPHOS maps, contrary to existing literature reporting greater glucose uptake of functional hubs [78].

### Energy pathways track developmental milestones

Energy metabolic pathways are tightly coupled to nutrient availability and exhibit adaptive changes across the lifespan, underlying their integral role in supporting neurogenesis and synaptic growth and integrity. To investigate the developmental trajectory of energy pathways, we used the BrainSpan RNA-sequencing data [79]. As before, energy pathway gene sets were used to retrieve pathway-specific sample-by-gene expression matrices and expression was averaged across genes. Given the importance of ketone bodies as an alternative energy substrate during early development, we also looked into the expression trajectory of genes involved in ketone body utilization (Table S4).

Fig. 6 illustrates the distinct trajectories of these energy pathways. Glycolysis, TCA, OXPHOS, and lactate metabolism exhibit a similar trajectory, rising from the fetal stage to infancy, with OXPHOS peaking in childhood and subsequently showing a decline into the adolescent and adult levels. This is in accordance with previous reports of glucose and oxygen uptake peaking in childhood, followed by a decrease in oxidative pathway activity by adolescence [12, 19]. This trend further resembles the trajectory of synapse development genes (Fig. S5). The PPP pathway shows a sharp decline from the fetal stage to infancy, followed by a gradual decrease into adolescence and adulthood. This underscores the importance of the PPP in supporting brain tissue generation during early development [80–82] and closely aligns with the lifespan trajectory of neural progenitor cells (Fig. S5). Furthermore, we find that the average expression of genes involved in ketone body utilization increases from the fetal stage to infancy, before declining in early childhood, consistent with the postnatal trajectory of ketone body utilization during the nursing and weaning periods [4, 19, 83]. Collectively, these findings reveal the unique expression trajectories of energy metabolic pathways across the lifespan, offering complementary insight into the metabolic dynamics shaped by developmental needs and nutrient availability.

**Figure 6.**
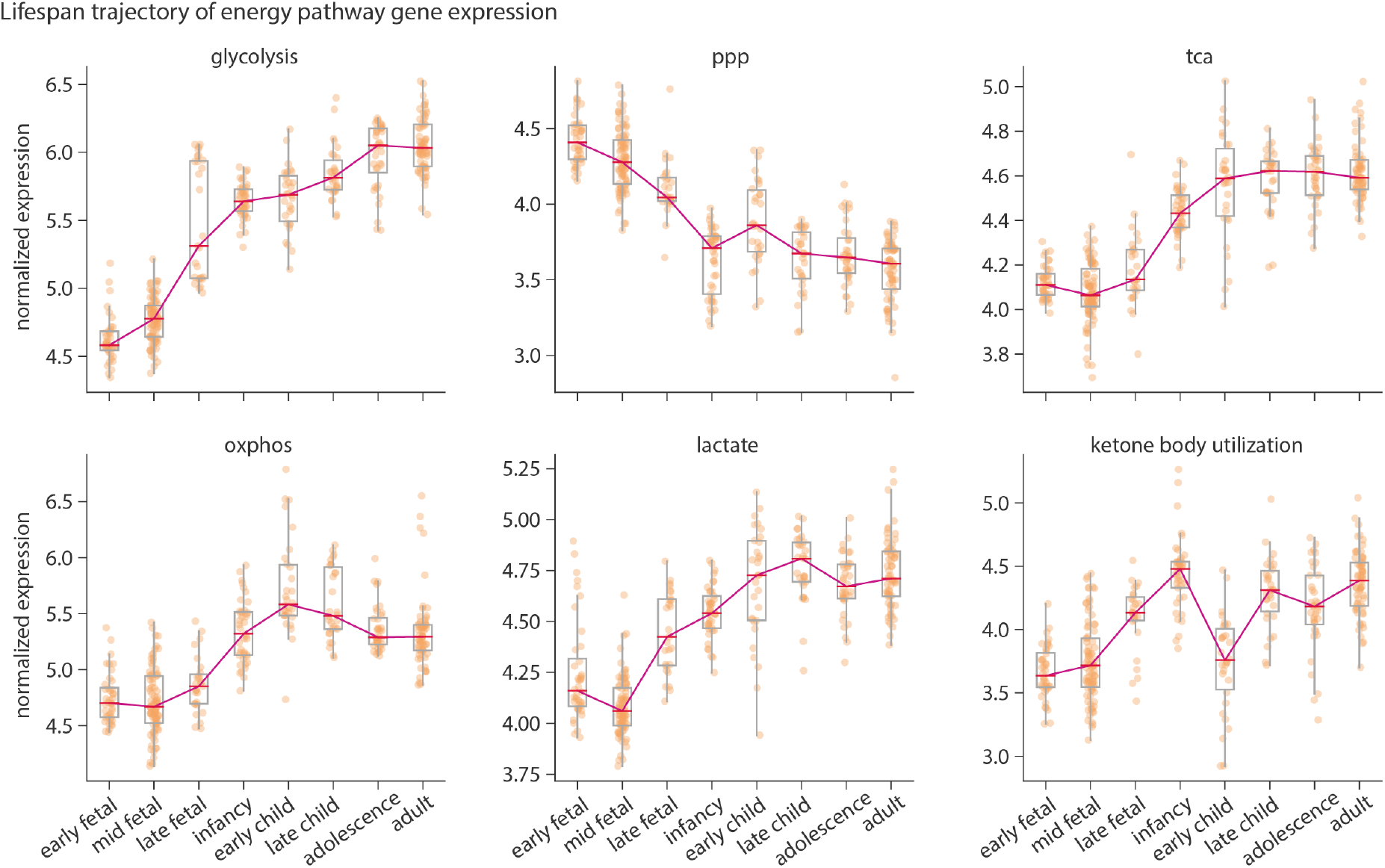
Lifespan trajectory of energy pathway gene expression. Developmental trajectories of energy metabolism pathways were produced using the BrainSpan RNA-sequencing dataset [79]. For each energy pathway, mean expression was calculated across all genes for each sample. Median expression across all samples falling into each age group was then calculated. Analysis only included cortical regions. The y-axis represents normalized log_2_(RPKM) expression values (see *Methods*. Dots represent samples in each age group. Line plot depicts the trajectory of median gene expression across age groups. For details of ages included in each group see Table S3. PPP, pentose phosphate pathway; TCA, tricarboxylic acid cycle; OXPHOS, oxidative phosphorylation; lactate, lactate metabolism and transport.

### Sensitivity analysis

As a final step, to ensure that the results are not influenced by preprocessing and analytic choices, we employ several sensitivity measures. First, given that only two out of six donors in the AHBA include samples from the right hemisphere, we reproduce energy maps using expression data exclusively from the left hemisphere. Maps derived from the left hemisphere correlate significantly with those produced by mirroring across hemispheres (Fig. S6, top panel). Second, we reproduce the maps and repeat the analyses with the lower resolution Schaefer-100 parcellation. The spatial distribution of energy pathways remains consistent with the original analysis (Fig. S6, bottom panel). Third, in order to analyze the lifespan trajectories independent of age categorization, we repeat the analysis using the non-parametric LOESS method against log(age) (Fig. S7). Lastly, we replicate the lifespan analysis using the BrainSpan microarray dataset [84]. Overall, the lifespan trajectories remain consistent with the original results (Fig. S8).

Finally, in addition to these five maps of key energy pathways, we also produced a complementary set of maps corresponding to other energy-related metabolic processes in the brain. These extended maps include: individual maps for the five mitochondrial complexes, ketone body utilization, fatty acid metabolism, glycogen metabolism, pyruvate dehydrogenase complex (PDC; responsible for the entry of pyruvate to the TCA cycle), malate-aspartate shuttle (MAS), glycerol phosphate shuttle (GPS), creatine kinase activity, detoxification of reactive oxygen species (ROS detox), generation of reactive oxygen species (ROS gen) nitric oxide signaling via guanylate cyclase (associated with vasodilation), Na^+^/K^+^ ATPase pump and the glutamine-glutamate cycle,(Fig. S9). For the lifespan trajectories of these extended energy processes see Fig. S12. Notably, mitochondrial ETC complexes show strong alignment with each other both in their spatial distribution across the cortex (rh̃o = 0.68, Fig. S11) and their lifespan trajectories (Fig. S12), reflecting their structural and functional coupling within the ETC. The greatest correlations are observed between complex I, III and IV, which reportedly form the most common supercomplexes. In contrast, complex II shows weaker correlations with the other complexes, which can be attributed to its less frequent incorporation in supercomplex formation, as well as involvement in TCA [85–87].

## DISCUSSION

In this study, we use a gene expression atlas to produce maps of energy metabolism pathways in the human brain. We explore the spatial distribution of five key energy pathways across the cortex and their correspondence to the structural and functional organization of the human brain at the micro- and macro-scale. We find that spatial patterns of energy pathway expression align with aspects of the information processing hierarchy. We observe a consistent dichotomy between energy producing pathways and pathways involved in cellular anabolic processes. Furthermore, the developmental trajectories of these pathways follow critical milestones, reflecting shifts in metabolic requirements and tissue generation capacity across the lifespan. Finally, we replicate our results using alternative processing approaches and datasets to ensure the robustness of these findings. This study leverages pathway gene expression as a complementary approach to elucidate the biochemical backbone of energy metabolism in the brain.

We find that energy pathways show heterogeneous gene expression across the cortex and are differently enriched across structural and functional networks. Pathways mainly involved in energy production and oxidative metabolism (glycolysis and OXPHOS) show a similar expression profile. This is consistent with the literature suggesting that the brain mainly relies on aerobic respiration to generate ATP [5, 6, 10, 88]. The glycolysis and OXPHOS maps show higher expression in the primary motor cortices and lower expression in the visual cortex, potentially hinting at differences in baseline energy requirements between these regions. The primary motor cortex, which generates efferent signals via large pyramidal neurons (Betz cells), demands substantial energy to support long-range axonal projections. On the other hand, the visual cortex is specialized for processing sensory input, has mostly short-range projections, and may rely on efficient encoding strategies and energy use. Interestingly, the creatine kinase (CK) map shows greater expression in the visual cortex (Fig. S9). CK catalyzes the reversible interconversion of creatine and phosphocreatine by transferring the phosphate group between ATP and ADP. The creatine-phosphocreatine shuttle therefore allows for fast regeneration of the ATP pool by continuous delivery of phosphate. CK is highly expressed in cells with high energy demand and is associated with sites of ATP production and consumption, such as mitochondria and ATPases (i.e. the sodium/potassium pump) [89, 90]. The higher expression of CK, together with the greater expression of ATPase pump components (Fig. S9), in visual cortex potentially allows for a rapidly mobilizable pool of ATP, coupling substrate oxidation to the creatine-phosphocreatine shuttle and bypassing the electron transport chain, while also reducing ROS production in mitochondria and exerting an indirect antioxidant effect [89].

The PPP displays a distinct spatial pattern, with greater expression in primary sensory cortices. The PPP is a parallel pathway to glycolysis and oxidative metabolism, that is involved in tissue building anabolic processes and antioxidant defense. It generates NADPH to maintain glutathione in a reduced state to neutralize reactive oxygen species, and produces ribose-5-phosphate for nucleotide and amino acid synthesis [13, 19, 91]. The higher expression of genes involved in the PPP in primary sensory cortices may potentially reflect the greater demand for accuracy and robustness in sensory encoding [47], necessitating active maintenance of synaptic integrity through ongoing cellular biosynthesis. The PPP map also aligns with gene PC1 gradient and the T1w/T2w ratio maps (Fig. S13). The T1w/T2w map has been proposed as an index of hierarchical organization in the human brain that aligns well with both tract-tracing models and captures the cyto- and myelo-architecture boundaries [92]. The gene PC1 gradient represents the principal axis of transcriptional variation across the cortex, reflecting fundamental differences in microscale organization that follows the cortical hierarchical organization, differentiating the primary sensory and motor cortices from higher order association areas. This is in concordance with the PPP’s role in tissue building and synthesis of fatty acids and cholesterol —essential for myelin production— and further underscores PPP as a fundamental component of brain microstructure and organization [81, 93, 94]. The TCA map aligns with both glycolysis/OXPHOS maps and the PPP map, signifying both its role as the link between glycolysis and OXPHOS, and involvement in cellular anabolic processes. These spatial relationships are also reflected in the correspondence between energy maps and the first and second FC gradients. FC gradients represent the dominant differences in connectivity patterns across the cortex, which recapitulates the unimodal-transmodal hierarchy [48]. Glycolysis and OXPHOS align with the second functional gradient, which differentiates between the unimodal cortices. In contrast, the PPP and TCA pathways capture the cortical hierarchy defined by the first functional gradient, separating primary regions from higher-order association and limbic cortices.

To validate the gene expression-based energy maps, we compared them to average maps from *in vivo* PET imaging of metabolism. Surprisingly, we find that the glycolysis and OXPHOS maps do not overlap significantly with the resting-state glucose and oxygen consumption or cerebral blood flow measured by PET. In contrast, the PPP shows the greatest correspondence with the CMR_*O*2_ map. A recent study has shown that the PPP metabolites exhibit the highest fold change, compared to glycolysis and TCA metabolites, in response to oxygen supplementation in newborn mice [95]. In addition to redox homeostasis, this may indicate biosynthetic demands for cellular repair in regions with greater oxygen consumption are consequently more susceptible to oxidative damage. The lactate map correlates most positively with the CMR_*glc*_ and GI PET maps, in line with lactate’s contribution to non-oxidative glucose use in the brain and the astrocyte-neuron lactate shuttle [9, 10, 96]. However, individual genes within these energy pathways exhibit a wide range of correlations with CMR_*glc*_, emphasizing the existence of multiple intermediate steps between gene transcription and pathway activity. Additionally, as PET imaging measures glucose uptake, the first step catalyzed by hexokinase, rather than downstream metabolic pathways, it is perhaps expected that it does not necessarily correlate with these downstream energy metabolism pathways.

Electrical activity in the brain relies on energy metabolism. Action potentials, synaptic transmission, and maintenance of membrane electrochemical gradients are energy-demanding processes critical for sustaining neuronal excitability and function [5, 97]. We therefore investigated the spatial overlap between MEG oscillatory activity and energy pathways. We find that glycolysis, TCA and lactate maps are most strongly associated with the MEG beta band. Beta oscillatory activity is associated with task activation and motor processes and is thought to arise from activity of GABAergic interneurons and bursting pyramidal cells [98]. The three energy pathways overlapping with beta activity comprise the astrocyte-neuron lactate shuttle, involved in rapid energy supply via transfer of lactate to neurons [10, 96, 99].

Different cell types in the brain have distinct yet complementary metabolic gene expression and enzyme activity profiles [100, 101]. To explore this organizational feature, we examined the relationship between energy maps and the distribution of seven major cell types in the cortex. Excitatory neurons show significant spatial alignment with the glycolysis and OXPHOS maps, in line with reports that glutamatergic signaling accounts for the majority of energy expenditure in the brain [3, 5, 102]. GABAergic interneurons are also reported to impose an energetic load on the brain, due to their high-frequency spiking and evidenced by large numbers of mitochondria in these cells [103–106]. Notably, we also find that inhibitory neurons show significant spatial overlap with all energy pathways. Reports suggest that inhibitory neurons are particularly vulnerable to oxidative stress [107, 108]. The alignment with lactate metabolism and PPP can therefore possibly hint at antioxidant defenses and molecular repair mechanisms in these cells. Lactate dehydrogenases catalyze the pyruvate:lactate interconversion and play an integral role in redox signaling and maintaining the NAD^+^:NADH balance within the cell [10, 109]. Furthermore, the PPP can provide bio-molecules required for cellular integrity and repair, as the greater presence of mitochondria and oxidative metabolism increases cellular damage. These results therefore point to both higher energy demands and antioxidant defenses in inhibitory neurons.

We also find that the microglia map correlates significantly with the OXPHOS map. It has been reported that microglia express genes for both glycolytic and oxidative pathways and mainly employ oxidative phosphorylation to meet their energy requirements [2, 71]. Furthermore, we find that endothelia correspond to the glycolysis and lactate maps, consistent with previous research suggesting that brain endothelial cell lines primarily engage in glycolytic metabolism *in vitro* [69]. The astrocyte map however, shows a negative correlation to the PPP and no correlation with the lactate map, contrary to the high PPP activity in astrocytes and their role in lactate production [73, 74, 96, 110]. It is important to note that metabolic compartmentalization can also occur at the protein level via post-translational modifications [9], which are not captured at the transcript level.

The hierarchical organization of brain function is also reflected in the cortical laminar architecture. This architecture is characterized by distinct cellular composition and specialized input-output circuitry, differences that likely impose specific energy demands to support their unique roles in information processing and integration [75, 111–113]. The PPP, TCA and lactate maps exhibit significant correlations with cortical layers 3 and 4. Granular layer 4 is the main input layer of the cortex, characterized by high vascular density [114, 115]. These results could therefore suggest mechanisms present for rapid energy supply upon the influx of sensory information and biosynthetic demands for accurate sensory encoding. Glycolysis and OXPHOS maps show significant alignment with the infragranular layer 5, the main output layer of the cortex. This potentially reflects the greater energy requirements of large pyramidal neurons such as Betz cells [75, 76, 111, 116].

We examined the relationship between the network properties of the cortex and energy pathways. Previous studies have reported greater glucose uptake in high-degree nodes within the functional connectivity network [55, 78]. In contrast to this, we find that node degree shows no correlation with glycolysis or OXPHOS gene expression maps. This may however indicate energy efficiency in these hub regions, as the relationship between glucose uptake and FC degree appears to be nonlinear [78], suggesting efficient glucose utilization. Functional connectivity strength shows a moderate overlap with the PPP pathway. This aligns with the theory that hub regions require elevated anabolic activity to support neuronal plasticity [66, 117, 118].

We also explored how energy metabolic pathways change across the human lifespan. Pathways primarily involved in ATP production, including glycolysis, TCA, OXPHOS and lactate metabolism, show a marked increase in expression from the fetal stage to infancy. This aligns with research indicating that cerebral glucose and oxygen consumption, initially low at birth, rise rapidly postnatally [19, 119, 120]. Notably, the mean OXPHOS expression decreases in adolescence and adulthood, in alignment with reports that oxygen consumption and the activity oxidative pathways decline by adolescence [19, 121, 122]. This pattern is overall consistent across individual mitochondrial complexes (Fig. S12) and also resembles the previously reported normative trajectory of total cortical and grey matter volume, which peaks in childhood [123–126] Additionally, genes involved in ROS detoxification exhibit reduced expression into adulthood (Fig. S12). This could reflect diminished cellular defenses against oxidative stress in later stages of life [127]. The PPP shows a postnatal decrease, followed by a gradual decline in later life, consistent with its integral role in fetal brain development and tissue generation and with previous reports on PPP enzymatic activity [12, 19, 81, 128, 129]. It has been demonstrated that the PPP activity progressively declines with age and is undetectable in the rat brain by 18 months of age [129]. Furthermore, PPP is one of the major contributors to aerobic glycolysis. Aerobic glycolysis is considered a measure of non-oxidative glucose metabolism in the presence of oxygen [25, 54, 130, 131] which is important for tissue biosynthesis [66, 132]. Previous reports of decreased aerobic glycolysis in the aging brain may therefore be attributed to diminished PPP activity [133]. The PPP trajectory also resembles the rate of growth of the mean cortical thickness across the lifespan, as well as the trajectories of cell proliferation genes and neural progenitor cells (Fig. S5) [79, 84, 123]. Collectively, these findings underscore the PPP’s critical role in providing anabolic support for brain tissue generation, emphasizing its importance for neurogenesis during fetal development [134] and highlighting reduced plasticity in the aging brain [129, 133, 135]. Additionally, given the PPP’s role in anti-oxidant defense and repair, this decline could further indicate reduced ROS-buffering capacity and increased susceptibility to oxidative stress in later life [127, 136, 137].

We also investigated pathways of ketone body metabolism. In the adult brain, ketone bodies are only a source of fuel during starvation, but in the infant brain they serve a critical role both as an energy source and a substrate for synthesis of cholesterol and other brain lipids [4, 83]. During the nursing period, the high fat content of maternal milk leads to elevated plasma concentrations of ketone bodies, making it an obligate fuel for the infant brain [138]. As weaning progresses and circulating ketone body levels decline, the brain shifts to glucose as fuel [19, 122, 139, 140]. Here, we demonstrate that this distinct post-natal pattern is also evident in the expression of genes involved in ketone body utilization. Mean cortical expression of ketone body utilization rises sharply from the fetal stage to infancy and declines in early childhood, mirroring the respective availability and use of ketone bodies during these developmental stages (Fig. 6, Fig. S8). The subsequent increase in late childhood could be associated with adiposity increase at this stage [141, 142] or point to a ketogenic shift in later life [143–145].

The present findings should be interpreted in light of several important limitations. First, the gene expression data used in this study were derived from only six post-mortem brains, affecting the generalizability of the results. Additionally, some genes involved in energy metabolic pathways were excluded based on quality control criteria. Furthermore, the AHBA samples were obtained at the adult stage (ages 24–57, 42.50 *±* 13.38) when the expression of energy metabolism genes and specifically the PPP genes has already declined. Nonetheless, the Allen Human Brain Atlas remains the most comprehensive transcriptomic atlas of the human brain, and we show that the results were consistent when restricting the analysis to the more extensively-sampled left hemisphere and when recreating the maps at a lower-resolution parcellation. Second, energy metabolism is characterized by interconnected pathways that branch out and converge via shared enzymes and metabolites [9]. Studying these pathways in isolation and relying on average expression measures is an oversimplification of the complex dynamics underlying energy metabolism. Third, transcript levels do not directly correspond to pathway activity, due to intervening regulatory steps (e.g., transcript splicing and stability, regulation of translation, post-translational modifications, protein ubiquitination, phosphorylation and degradation). Further advancements are needed to facilitate high resolution *in vivo* studies of energy pathways. Fourth, data used in the lifespan analyses is obtained from a different cohort and constitutes a limited number of samples at each developmental stage, affecting the generalizabitlity of the results. However, we show that the results remain consistent using a replication dataset.

In summary, we demonstrate the heterogeneous expression of key components of energy metabolism in the human brain. We show that these energy pathways show distinct alignment with the structural and functional organization of the cortex and exhibit dynamic trajectories across the lifespan. Collectively these results provide a complementary perspective on the complexity and organization of brain energy metabolism. The energy pathway maps, along with the data and scripts used in this study, are made publicly available to hopefully facilitate future research on brain metabolism.

## METHODS

### Energy pathway gene sets

Over the last decade, there has been an immense global effort to curate databases of the complex network of biological pathways in the living organisms. Databases such as Gene Ontology (GO) [39] and Reactome Knowledge base [40] are continuously updated and expanded to include evidence-based and cross-referenced research findings on molecular functions and biological pathways within a cell. These databases provide a unified platform for gene functional classification and pathway annotation and facilitate the study of biological systems, including genes, proteins and their interactions within a living organism. Here, we used the Gene Ontolgy and Reactome databases to curate gene sets pertaining to energy metabolism pathways. Pathways included in the main analysis include: glycolysis, pentose phosphate pathway (PPP), tricaboxylic acid cycle (TCA), oxidative phosphorylation (OXPHOS) and lactate transport and metabolism. For each of these pathways, we identified the corresponding GO biological processes [39] and Reactome pathway [40] IDs. We then retrieved gene sets involved in each pathways using the *biomaRt* version 2.50.3 ([146], https://bioconductor.org/packages/elease/bioc/html/biomaRt.html/ and *GO*.*db* packages (https://bioconductor.org/packages/GO.db/). *BioMart* is a freely available data-mining tool that provides unified access to biological knowledge bases. We used the Ensemble human gene annotation database release 112 [147] to retrieve gene sets for each pathway ID. For each pathway, genes consistently annotated in both databases were retained for further analysis. Of the three hexokinase enzymes, only hexokinase 2 met the differential stability criteria. However, since hexokinase catalyzes the first step of both glycolysis and the PPP and therefore entrance to either pathway, it was excluded from these gene sets.

To provide a more comprehensive view of energy metabolism in the brain, we also produced an extended set of maps including: individual maps for the five mitochondrial complexes, ketone body utilization, fatty acid metabolism, glycogen metabolism, pyruvate dehydrogenase complex (PDC; responsible for the entry of pyruvate to the TCA cycle), malate-aspartate shuttle (MAS), glycerol phosphate shuttle (GPS), creatine kinase (CK), detoxification of reactive oxygen species (ROS detox), generation of reactive oxygen species (ROS gen), the glutamine-glutamate cycle, nitric oxide signaling and Na^+^/K^+^ ATPase pump. MAS, GPS, CK and the glutamine-glutamate cycle gene sets were curated based on existing literature [19, 90, 148–150].

### Microarray gene expression data

Regional microarray expression data were obtained from 6 post-mortem brains (1 female, ages 24–57, 42.50 *±* 13.38) provided by the Allen Human Brain Atlas (https://human.brain-map.org; Allen Human Brain Atlas white paper), [38]. Data were processed using a 400-region volumetric atlas in MNI space. Microarray probes were reannotated using data provided by [151]; probes not matched to a valid Entrez ID were discarded. Next, probes were filtered based on their expression intensity relative to background noise [152], such that probes with intensity less than the background in *≥* 50% of samples across donors were discarded, yielding 31 569 probes. When multiple probes indexed the expression of the same gene, we selected and used the probe with the most consistent pattern of regional variation across donors (i.e., differential stability; [153].

MNI coordinates of tissue samples were updated to those generated via non-linear registration using the Advanced Normalization Tools (ANTs; https://github.com/ chrisfilo/alleninf). To increase spatial coverage, tissue samples were mirrored bilaterally across the left and right hemispheres [154]. Samples were assigned to brain regions in the provided atlas if their MNI coordinates were within 2 mm of a given parcel. If a brain region was not assigned a tissue sample based on the above procedure, every voxel in the region was mapped to the nearest tissue sample from the donor in order to generate a dense, interpolated expression map. The average of these expression values was taken across all voxels in the region, weighted by the distance between each voxel and the sample mapped to it, in order to obtain an estimate of the parcellated expression values for the missing region. All tissue samples not assigned to a brain region in the provided atlas were discarded. Inter-subject variation was addressed by normalizing tissue sample expression values across genes using a robust sigmoid function [155]. Normalized expression values were then rescaled to the unit interval [41]. Gene expression values were then normalized across tissue samples using an identical procedure. Samples assigned to the same brain region were averaged separately for each donor, yielding a regional expression matrix for each donor with 400 rows, corresponding to the cortical regions in the Schaefer-400 parcellation, and 15 633 columns, corresponding to the retained genes. From this initial expression matrix, we retained genes with a differential stability value greater than 0.1 [92], yielding expression data for a total of 8 687 genes.

Energy pathway gene sets were used to extract pathway-specific gene expression matrices. The final number of genes per pathway differs from the original sets, as some genes were not present in the AHBA dataset or were excluded based on the above-mentioned quality control criteria applied during preprocessing. the differential stability distribution for energy pathway genes is shown in (Fig. S14). For each pathway, expression values were averaged across all genes to yield a pathway mean gene expression map. Brain maps were plotted using the *surfplot* package [156, 157].

### Spatial auto-correlation preserving nulls

The brain is spatially constrained within the skull and exhibits inherent spatial auto-correlation in both structure and function. Data mapped onto the brain such as gene expression are not independent and identically distributed (i.i.d.), which is a common prerequisite for many statistical tests. Due to the spatial autocorrelation present in the brain, nearby voxels/regions are more likely to have similar values (e.g. similar gene expression) due to both biological (e.g., local connectivity) and technical (e.g., image processing and smoothing) factors [158]. This can lead to inflated false-positive rates in statistical testing of two or more brain maps.

To account for this, various spatial permutation tests (spin tests) have been introduced [159– Spin tests account for the spatial auto-correlation present in brain data by permuting voxels/parcels while maintaining the spatial structure and auto-correlation, therefore providing a more accurate framework for hypothesis testing. In this study, we used the spatial permutation test developed by Váša et al. [160] implemented in the netneurotools package (https://netneurotools.readthedocs.io/). This method uses parcel centroid coordinates to produce rotations and ensure that there are no duplicate reassignments, leading to a true null distribution. We refer to the non-parametric p-value calculated using spatial permutation testing as *p*_spin_.

### Parcellations, structural classes and functional networks

We used a cortical parcellation developed by Schaefer et al. [37] which divides the cortical surface into 400 regions. This parcellation was generated using a gradient-weighted Markov Random Field model from resting state fMRI data, integrating both local gradients and global similarity to define parcel boundaries. To investigate how our energy maps are distributed across functional networks, we used the resting state network assignments provided by the original authors which is based on the seven intrinsic functional networks described by Thomas Yeo et al. [52]. To explore networks pertaining to structural classes, we used two network definitions: von Economo-Koskinas cytoarchitectonic classes based on the morphology and laminar differentiation of neuronal types [49, 162, 163], and Mesulam classes describing the sensory-fugal hierarchy of information processing [47, 164].

For analysis of energy expression within the visual cortex, we used the Glasser parcellation ([53], https://github.com/brainspaces/glasser360/). Functional delineation of the visual cortex hierarchy was obtained from Glasser et al. [53] supplementary neuroanatomical results and https://neuroimaging-core-docs.readthedocs.io/en/latest/pages/atlases.html/. Subcortical energy maps were produced using the Desikan-Killiany atlas [165] and plotted using the ENIGMA Toolbox [166].

### Functional connectivity gradients

The first two functional connectivity (FC) gradients calculated by Margulies et al. [48] were retrieved from the *neuromaps* package [167]. Briefly, gradients were calculated for 820 healthy individuals from the Human Connectome Project (HCP) S900 release. The affinity matrix was then calculated from the FC matrix using cosine distance. FC gradients were computed using diffusion embedding, a non-linear dimensionality reduction technique that projects the data into a low-dimensional embedding space and assures a more stable representation of the connections compared to other dimensionality reduction techniques [48]. FC gradients were parcellated according to the Schaefer-400 atlas [37].

### Metabolic PET neuroimaging data

Metabolic PET maps were produced previously in [54] and retrieved from the *neuromaps* package [167]. These PET maps include: CMR_*glc*_ using [^18^*F*]-labeled fluorodeoxyglucose (FDG) radiotracer, CMR_*O*2_ and cerebral blood flow (CBF) using [^15^*O*]oxygen and water. All PET maps were obtained from neurologically normal individuals (*n* = 33, 14 males; age = 25.4 *±* 2.6 years) at resting state, using a Siemens model 961 ECAT EXACT HR 47 PET scanner [54]. A map of glycolytic index (GI) from the same study was also included in the analysis. The GI map is produced using the residuals after linearly regressing CMR_*glc*_ on CMR_*O*2_ and it was introduced previously as a measure of non-oxidative metabolism of glucose (aerobic glycolysis) [54]. Metabolic PET maps were then parcellated into the Schaefer-400 parcellation using the *neuromaps* package.

### Magnetoecephalography maps

MEG frequency data were first processed and used in [61] and were retrieved using the *neuromaps* package [167]. Resting state MEG data of a set of healthy young adults (n =33, 17 males; age 22–35 years) with no familial relationships were obtained from HCP (S900 release; [168]). The data include resting state scans of about 6 minutes duration (sampling rate =2 034.5 Hz; anti-aliasing lowpass filter at 400 Hz) and noise recordings for all participants. The data was analyzed using BrainStorm [169]. Pre-processing was performed by applying notch filters at 60, 120, 180, 240 and 300 Hz and was followed by a high-pass filter at 0.3 Hz to remove slow-wave and DC-offset artifacts. The artifacts (including heartbeats, eye blinks, saccades, muscle movements, and noisy segments) were then removed from the recordings using automatic procedures as proposed by Brainstorm. Pre-processed sensor-level data were used to obtain a source estimation on HCP’s fsLR4k cortex surface for each participant. Head models were computed using overlapping spheres, and the data and noise covariance matrices were estimated from the resting-state MEG and noise recordings. Brainstorm’s linearly constrained minimum variance beamformers method was applied to obtain the source activity for each participant. Data covariance regularization was performed using the “median eigenvalue” method from Brainstorm[169]. The estimated source variance was also normalized by the noise covariance matrix. Source orientations were constrained to be normal to the cortical surface at each of the 8004 vertex locations on the fsLR4k surface. Welch’s method was then applied to estimate power spectrum density for the source-level data, using overlapping windows of length 4 seconds with 50% overlap. Average power at each frequency band was then calculated for each vertex as the mean power across the frequency range of a given frequency band. The power spectrum was computed at the vertex level across six canonical frequency bands: delta (2–4 Hz), theta (5–7 Hz), alpha (8–12 Hz), beta (15–29 Hz), low gamma (30–59 Hz) and high gamma (60–90 Hz). Group-averaged maps for each MEG frequency bands were retrieved from the *neuromaps* package [167] and parcellated according to the Schaefer-400 cortical atlas [37].

### Cell and layer specific gene expression maps

Brain cell type and layer specific gene sets were obtained from Wagstyl et al. [62], supplementary file 2. Briefly, the authors curated cell-type markers by combining data from five single-cell and single-nucleus RNAsequencing studies [79, 170–176]. Subcategories across these studies were grouped into seven canonical classes including: excitatory neurons, inhibitory neurons, astrocytes, endothelial cells, microglia, oligodendrocytes and oligodendrocyte progenitor cells. Cell type gene sets were used to filter the AHBA gene expression matrix to obtain region-by-gene cell-specific expression matrices. Expression was then averaged across genes to yield a cell type mean expression map corresponding to the Schaefer-400 parcellation. Layer specific gene sets were curated based on two RNA-sequencing studies using samples from the prefrontal cortex [177, 178]. Layer-specific gene sets were unionized across these two studies by Wagstyl et al. [62]. Layer-specific mean expression maps were produced as above.

### Functional and structural connectivity measures

Both functional magnetic resonance imaging (fMRI) and diffusion weighted imaging (DWI) data were previously obtained for 326 unrelated participants (145 males; age 22–35 years) from the Human Connectome Project (HCP) S900 release [67, 168, 179]. fMRI data was acquired using a 3T scanner for 15 minutes during the resting state. All 4 resting state fMRI scans (2 scans with R/L and L/R phase encoding directions on day 1 and day 2, each about 15 minutes long; TR =720 ms) were available for all participants. Preprocessing was previously performed using the minimal preprocessing pipeline [180]. Briefly, all 3T functional MRI time-series (voxel resolution of 2mm isotropic) were corrected for gradient nonlinearity, head motion using a rigid body transformation, and geometric distortions using scan pairs with opposite phase encoding directions (R/L, L/R) [128]. Further preprocessing steps include coregistration of the corrected images to the T1w structural MR images, brain extraction, normalization of whole brain intensity, high-pass filtering (>2,000s FWHM; to correct for scanner drifts), and removing additional noise using the ICA-FIX process [67, 181]. The fMRI time series were then parcellated into 400 cortical regions in the Schaefer atlas [37] and the functional connectivity matrix was generated by computing the Pearson’s correlation coefficient between pairs of regional time series for each of the 4 scan each participant. A group-average functional connectivity matrix was computed representing mean functional connectivity across all subjects and normalized using a Fisher’s r-to-z transformation [182, 183].

Structural connectomes were previously generated from minimally processed HCP S900 DWI data using the MRtrix3 package [184]. Multi-shell and multitissue response functions were estimated and spherical-deconvolution informed filtering of tractograms (SIFT2) was applied to reconstruct whole brain streamlines weighted by cross-section multipliers [185, 186]. The initial tractogram was generated with 40 million streamlines, with a maximum tract length of 250. For each subject, these reconstructed cross-section streamlines were mapped onto the Schaefer-400 atlas [37] to build a structural connectome. A group-consensus binary network was constructed, preserving the density and edge-length distributions of the individual connectomes [187].

Graph theory can be used to study the brain as a network of interconnected nodes and edges, where nodes represent distinct regions or units within the brain (i.e., neurons or parcels), and edges are the connections or interactions between these units (structural connection or functional interactions between pairs of nodes) [188–190]. Graph theory therefore allows us to define measures of regional importance in the brain connectivity network. We used the *bctpy* package (https://github.com/aestrivex/bctpy) to calculate these network measures from the binary structural connectome and the functional connectivity matrix including degree centrality and strength [191]. Below, we will briefly describe each measure. **Degree** a node’s degree is defined as the number of edges (connections) of that node. It is a local network measure calculated at each node that represents “hubness” in a network. **Strength** is the weighted analogue of degree centrality. In a weighted matrix such as the functional connectivity matrix, strength is calculated as the sum of connection weights incident on a node (here average time series correlation, in the functional connectivity matrix).

### Human brain lifespan transcriptomics data

BrainSpan is a freely available database containing developmental transcriptomics of the human brain spanning the pre-natal stages to adulthood https://www.BrainSpan.org/static/download.html/. The data includes 524 samples from 42 donors across 31 developmental stages spanning from 8 weeks post conception (PCW) to 40 years of age. The samples were taken from a total of 26 cortical and sub-cortical regions in the brain (BrainSpan white paper). RNA-sequencing data was previously processed into normalized RPKM (Reads Per Kilobase of transcript per Million) values using conditional quantile normalization to account for GC content and sequencing depth and batch effect correction using ComBat [79, 192].

RNA-sequencing data (Genecode v10 summarized to genes) were downloaded from the BrainSpan database (BrainSpan Download). The expression matrix contains normalized RPKM values for 52 376 genes across 524 samples. We grouped the samples into major developmental stages including: early fetal, mid fetal, late fetal, infancy, early childhood, late childhood, adolescence and adulthood ([84], Table. S3). All subsequent analysis was performed on the cortical samples. First, We carried out a basic cleanup of the sample-by-gene expression matrix: (1) We retained regions that had at least 1 sample in each age group. (2) Duplicate genes were removed, yielding 47 808 unique genes. (3) Genes were retained if they had an RPKM value >= 1 in 80% of the samples at each spatiotemporal point [193].

The cleanup step resulted in a 352 samples (155 females) from 11 cortical regions and 8 370 genes. Cortical regions include: anterior (rostral) cingulate (medial prefrontal) cortex, dorsolateral prefrontal cortex, inferolateral temporal cortex (area TEv area 20), orbital frontal cortex, posterior (caudal) superior temporal cortex (area 22c), posteroventral (inferior) parietal cortex, primary auditory cortex (core), primary motor cortex (area M1 area 4), primary somatosensory cortex (area S1 areas 312), primary visual cortex (striate cortex area V1/17) and ventrolateral prefrontal cortex.

Expression values were log_2_ transformed and normalized using the upper quartile method [194–196]. Each donor’s data was scaled by their 75th percentile expression value and multiplied by the mean 75th percentile value across all donors. We then retrieved sample-bygene expression matrices for each energy pathway using curated gene sets. Average expression across all genes for each energy pathway was calculated for each sample. Mean energy expression was then aggregated into the eight age groups by combining all samples within each respective age group. Pathway expression was then plotted across age categories. Marker genes for neural progenitor cells and synapse development were obtained from the supplementary materials of Kang et al. [84] and Li et al. [79]. Smoothed curves were produced using the LOESS method against log_10_(age) in post conception days using the *rpy2* package https://rpy2.github.io/doc.html/. For the microarray dataset, genes were retained if they had a log_2_(expression) *≥* 6 [84] and the rest of the analysis was carried out as above.

### Data and code availability

All the scripts and data used to perform the analyses are available at https://github.com/netneurolab/pourmajidian_metabolism-genes/. Biological databases are openly accessible in *biomaRt* at https://bioconductor.org/packages/release/bioc/html/biomaRt.html/. The Allen Human Brain Atlas is available at https://human.brain-map.org/static/download/. The source data from the Human Connectome Project S900 release) including diffusion-weighted MRI, functional MRI and MEG are available at https://db.humanconnectome.org/. Group-averaged PET, MEG and functional connectivity gradient images are available in *neuromaps* at https://netneurolab.github.io/neuromaps/listofmaps.html/. BrainSpan RNA-sequencing and microarray datasets are available at https://www.brainspan.org/static/download.html. The Schaefer-400 and 100 parcellations are openly available at https://github.com/ThomasYeoLab/CBIG/tree/master/stable_projects/brain_parcellation/Schaefer2018_LocalGlobal/Parcellations/ and can also be retrieved using *abagen* at https://abagen.readthedocs.io/. The Glasser parcellation is available at https://github.com/brainspaces/glasser360.

## Acknowledgment

We thank Vincent Bazinet, Eric Ceballos, Asa Farahani, Zhen-Qi Liu, Andrea Luppi, Filip Milisav, Filip Morys and Andrew Vo for their comments and suggestions on the paper. AD acknowledges support from the Canadian Institutes of Health Research (CIHR) Foundation scheme. BM acknowledges support from the Natural Sciences and Engineering Research Council of Canada (NSERC), Canadian Institutes of Health Research (CIHR), Brain Canada Foundation Future Leaders Fund, the Canada Research Chairs Program, the Michael J. Fox Foundation, and the Healthy Brains, Healthy Lives initiative (HBHL). BM acknowledges support from the Molson foundation and the Healthy Brains, Healthy Lives. MP acknowledges support from the Healthy Brains, Healthy Lives initiative (HBHL) and the Harold and Audrey Fisher Training Studentship. JYH acknowledges support from the Helmholtz International BigBrain Analytics and Learning Laboratory and the Natural Sciences and Engineering Research Council of Canada. GS was supported by a postdoctoral fellowship from the Canadian Institutes of Health Research (CIHR). The funders had no role in study design, data collection and analysis, decision to publish or preparation of the manuscript.

---

**Figure S1.**
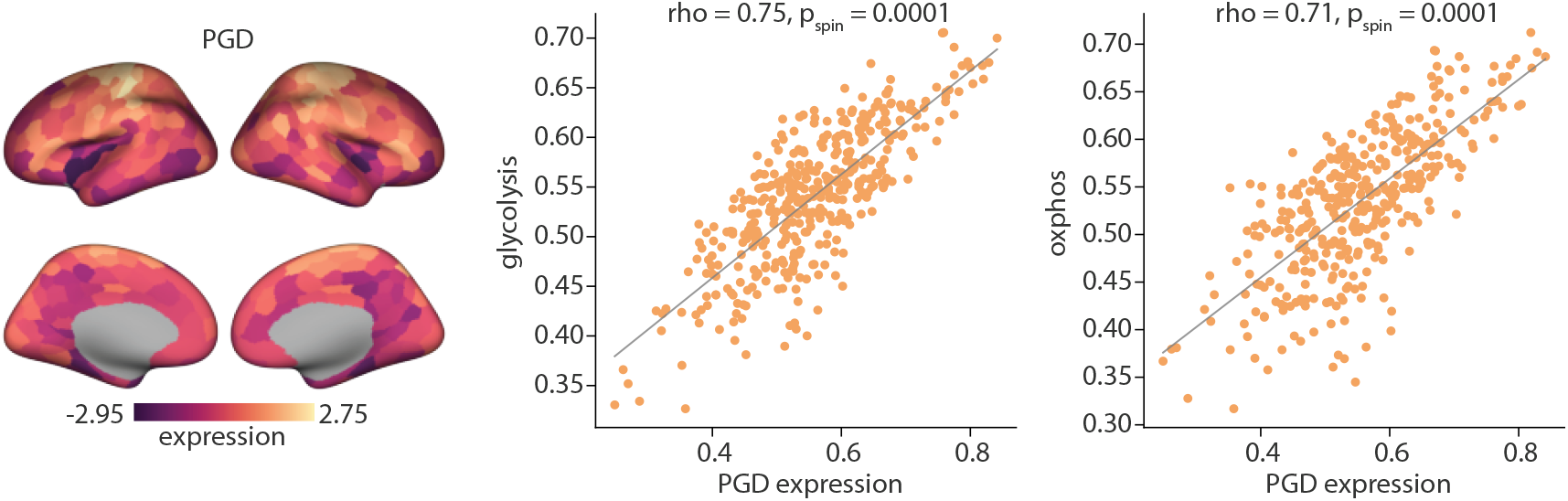
PGD gene expression correlates with glycolysis and OXPHOS maps. Brain map depicts PGD expression according to the Schaefer-400 parcellation. Color bar represents z-scored expression values. PGD expression was correlated (Spearman’s) with glycolysis and OXPHOS mean expression maps. Correlations were tested against a distribution of 10 000 correlations produced from the spatial permutation testing. The non-parametric p-value is indicated as *p*_spin_. Dots in the scatter plot represent 400 cortical regions in the Schaefer-400 parcellation.

**Figure S2.**
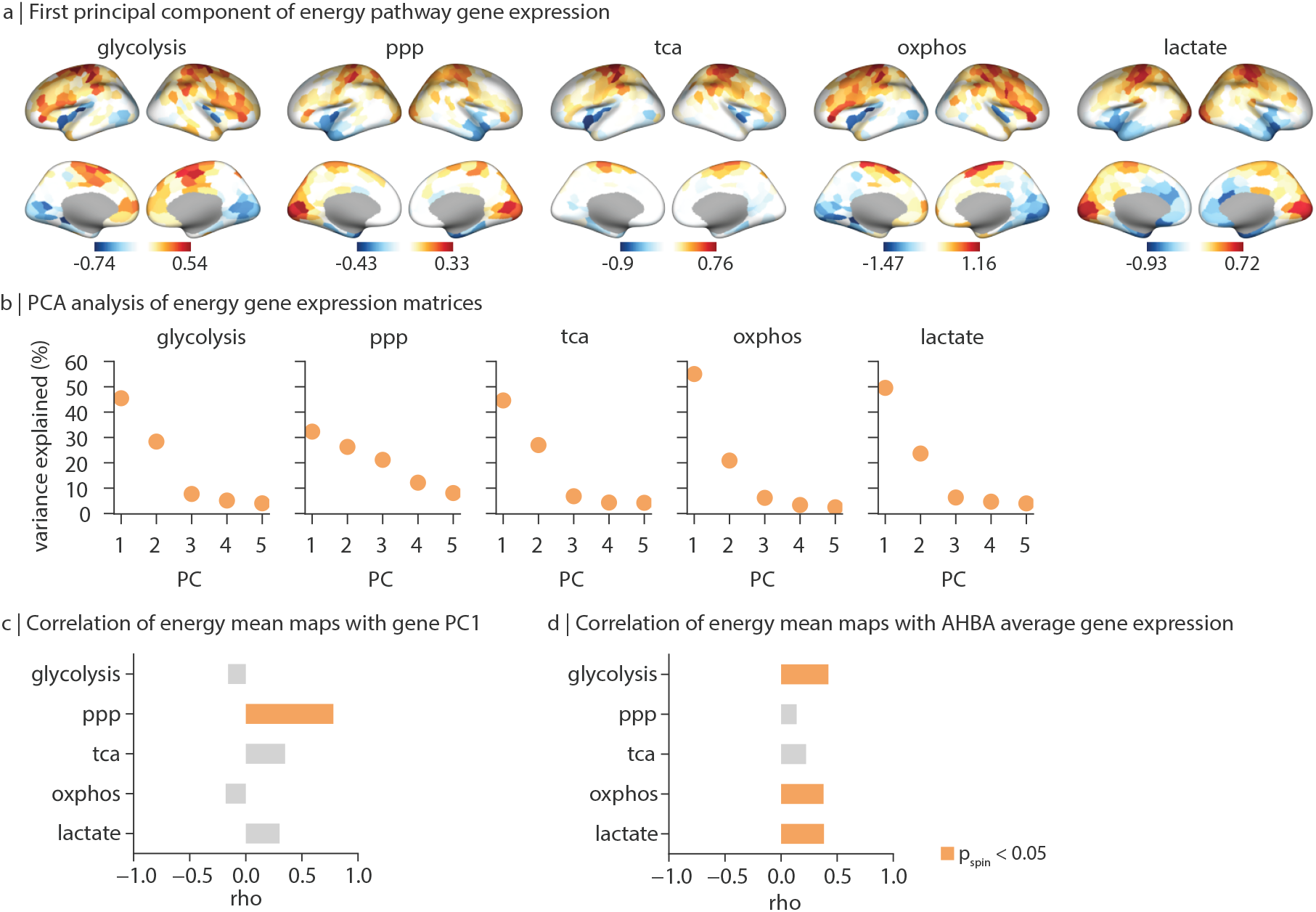
Principal component analysis of energy pathway gene expression. (a) Brain maps showing the first principal component of pathway gene expression matrices. PC1 of the glycolysis and OXPHOS gene expression reflect a gradient from the motor and prefrontal cortices to the parietal association regions, and finally the visual cortex (glycolysis: *var*_*explained*_ = %45.46; OXPHOS: *var*_*explained*_ = %55.01). PC1 of the PPP gene expression shows a spatial pattern closely capturing the established global gene expression gradient, extending from the sensory cortices to the higher order association, and limbic areas [92] (*var*_*explained*_ = %32.30, Fig. S2). (b) percent of variance explained by the first five principal components. (c) Spearman’s correlation between energy maps and the PC1 of all genes in the AHBA. (d) Correlation between energy maps and average expression of all genes in the AHBA. Highlighted bars represent statistical significance when tested against a distribution of 10 000 correlations produced from the spatial permutation testing (*p*_spin_ < 0.05). PPP, pentose phosphate pathway; TCA, tricarboxylic acid cycle; OXPHOS, oxidative phosphorylation; lactate, lactate metabolism and transport.

**Figure S3.**
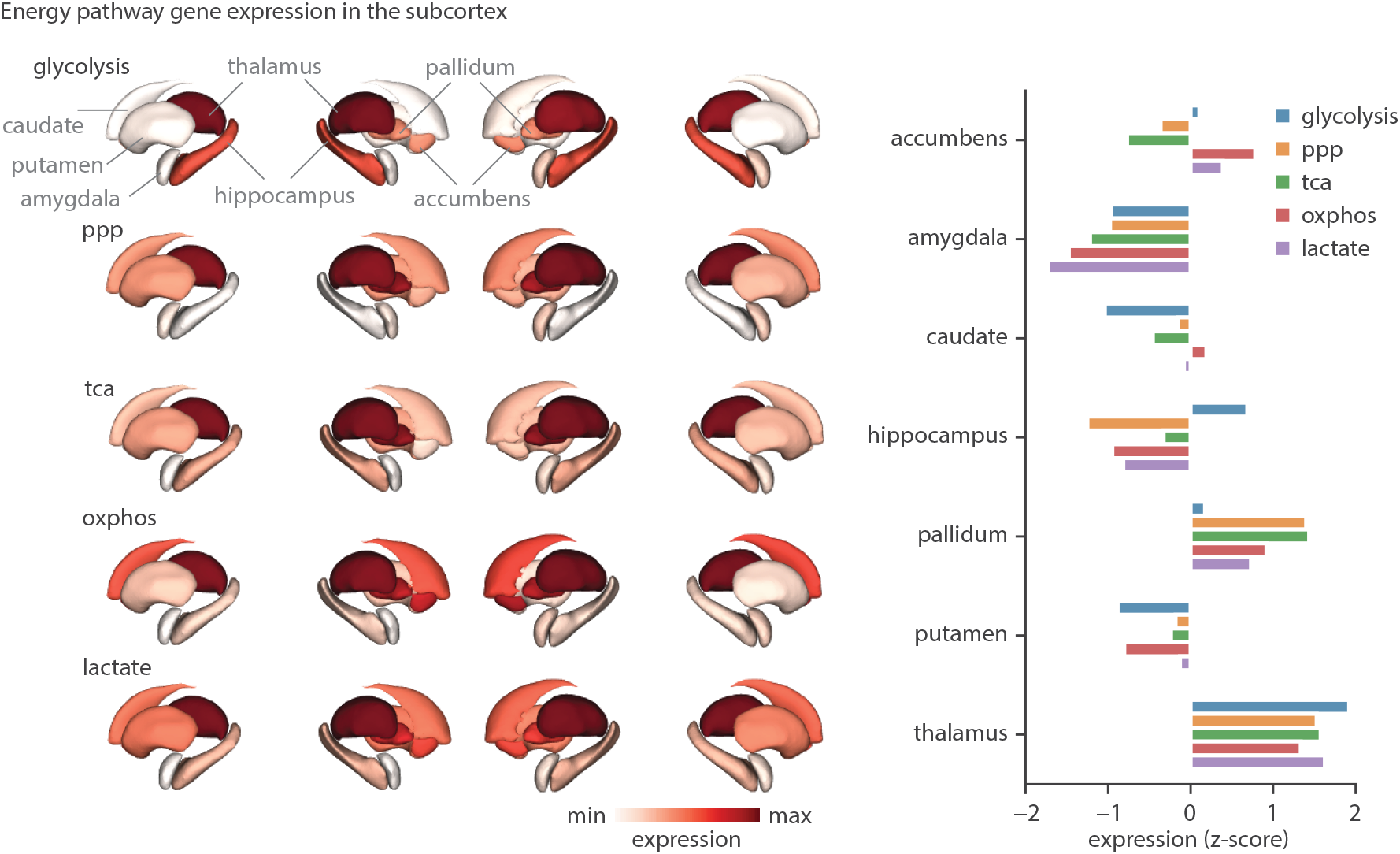
Subcortical energy pathway profiles. Energy pathway gene expression matrices were retrieved for 14 subcortical regions in the Desikian-Killiany (DK) atlas [165]. Left: subcortical visualization of mean pathway gene expression. Ventricles are excluded due to the absence of gene expression data. Stable genes (ds > 0.1) were retained to produce pathway mean gene expression maps (see *Methods*). Right: Barplot representation of the subcortical energy profiles (left hemisphere). Bars correspond to mean pathway gene expression, z-scored across all subcortical regions. Energy pathways consistently show higher expression in the thalamus and lower expression in the amygdala [197].

**Figure S4.**
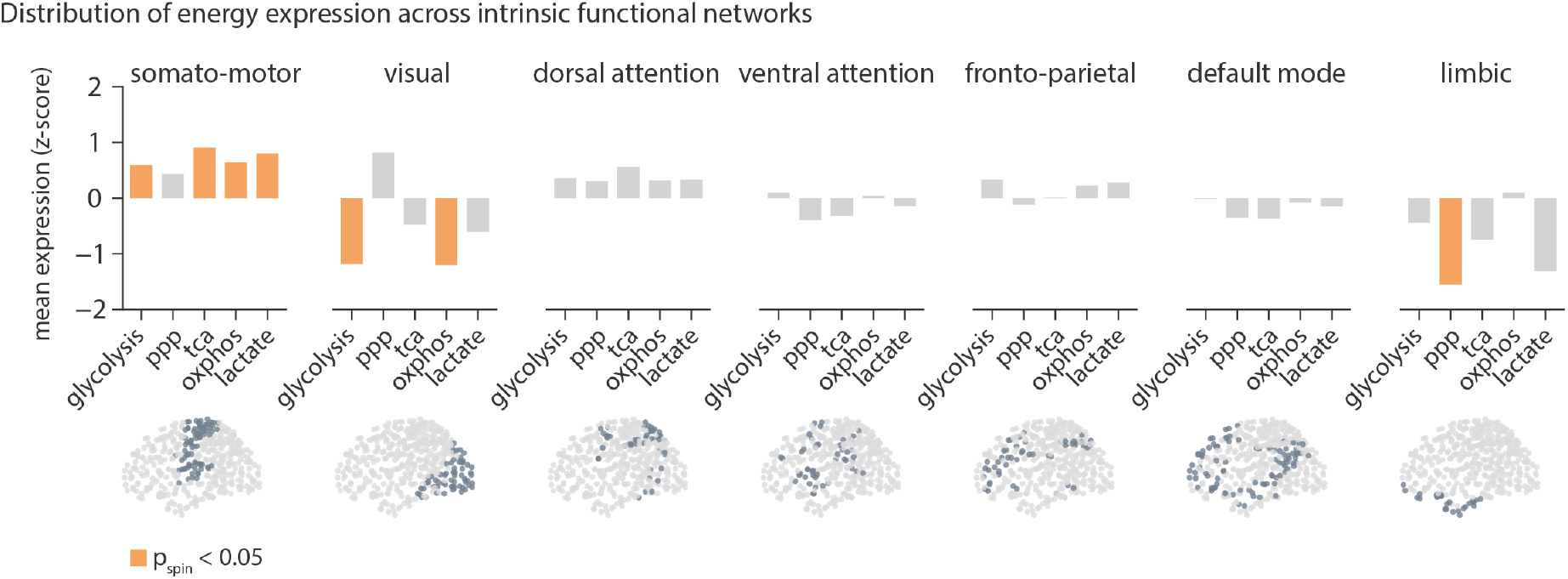
Distribution of energy pathway maps across intrinsic functional networks. Maps were z-scored across the 400 cortical regions and the average expression of parcels falling into each functional network was calculated for each energy map, according to the Yeo-Kiernen intrinsic functional network parcellation [52]. Highlighted bars indicate statistical significance when tested against 10 000 spatial-autocorrelation preserving nulls (*p*_spin_ < 0.05). Brain plots visualize each functional network. Glycolysis, TCA, OXPHOS and lactate maps show significantly greater values in the somato-motor cortex (glycolysis: *p*_spin_ = 0.04; TCA: *p*_spin_ = 0.00; OXPHOS: *p*_spin_ = 0.03; lactate: *p*_spin_ = 0.02). and significantly lower expressions in the visual cortex (glycolysis: *p*_spin_ = 0.00; OXPHOS: *p*_spin_ = 0.00). The PPP map on the other hand shows greater expression in the visual cortex, although not significant when tested against spatial permutations (*p*_spin_ = 0.09) and significantly lower expression in the limbic network (*p*_spin_ = 0.00)

**Figure S5.**
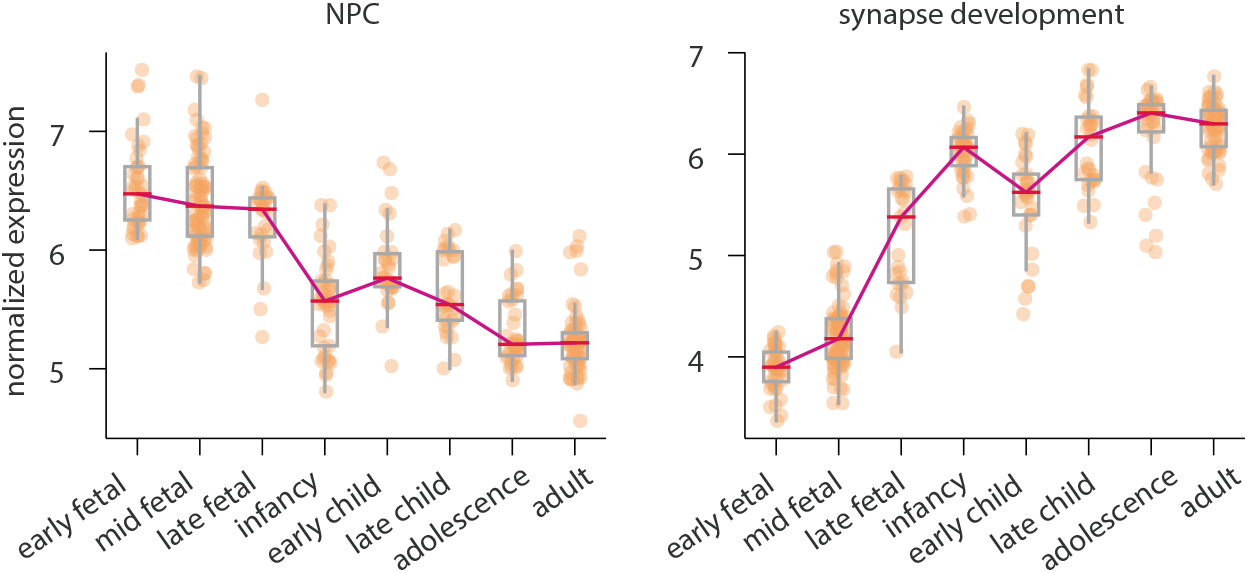
Expression trajectory of genes related to developmental processes. Lifespan trajectory of developmental processes were produced using previously curated gene sets [79, 84]. Mean expression of marker genes for each developmental process was calculated for each sample. Samples were then grouped into age categories. Line plot represents median values across all samples in each age group. Analysis only included cortical regions. The y-axis represents normalized log_2_(signal intensity) (see *Methods*). Dots represent individual samples in each age group. For details of ages included in each group see Table S5. NPC, neural progenitor cells.

**Figure S6.**
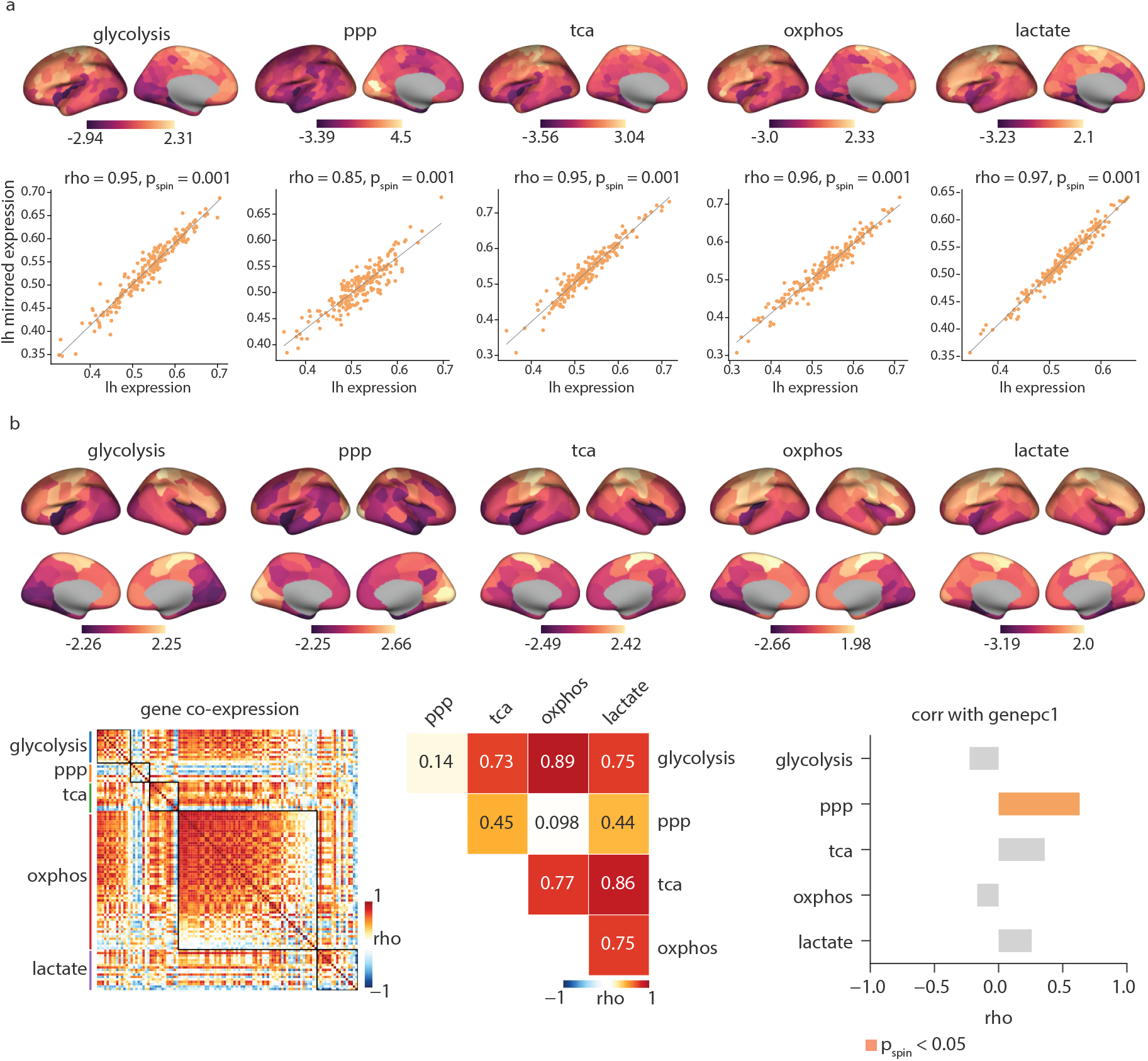
Sensitivity analysis. To assess the robustness of the results, energy maps were reproduced using (1) only the left hemisphere microarray data, and (2) a coarser parcellation (Schaefer-100). (a) Maps produced using only left hemisphere data and their correlation with the left hemisphere of original maps, produced by mirroring across hemispheres. Scatter plot represents 200 regions in the left hemisphere parcellated according to Schaefer-400 parcellation. (b) Energy maps according to Schaefer100 parcellation. Left: Heatmap depicts the pairwise correlation among all genes included in the energy sets across 100 regions in the Schaefer-100 parcellation. Middle: Spearman’s correlation between mean expression energy maps. Right: correlation of energy maps with the first principal component of all genes in AHBA (gene pc1). Brain map color bars represent z-scored expression across all regions in each parcellation. Highlighted bars show statistical significance when tested against 1 000 spatial-autocorrelation preserving nulls. lh, left hemisphere; PPP, pentose phosphate pathway; TCA, tricarboxylic acid cycle; OXPHOS, oxidative phosphorylation; lactate, lactate metabolism and transport.

**Figure S7.**
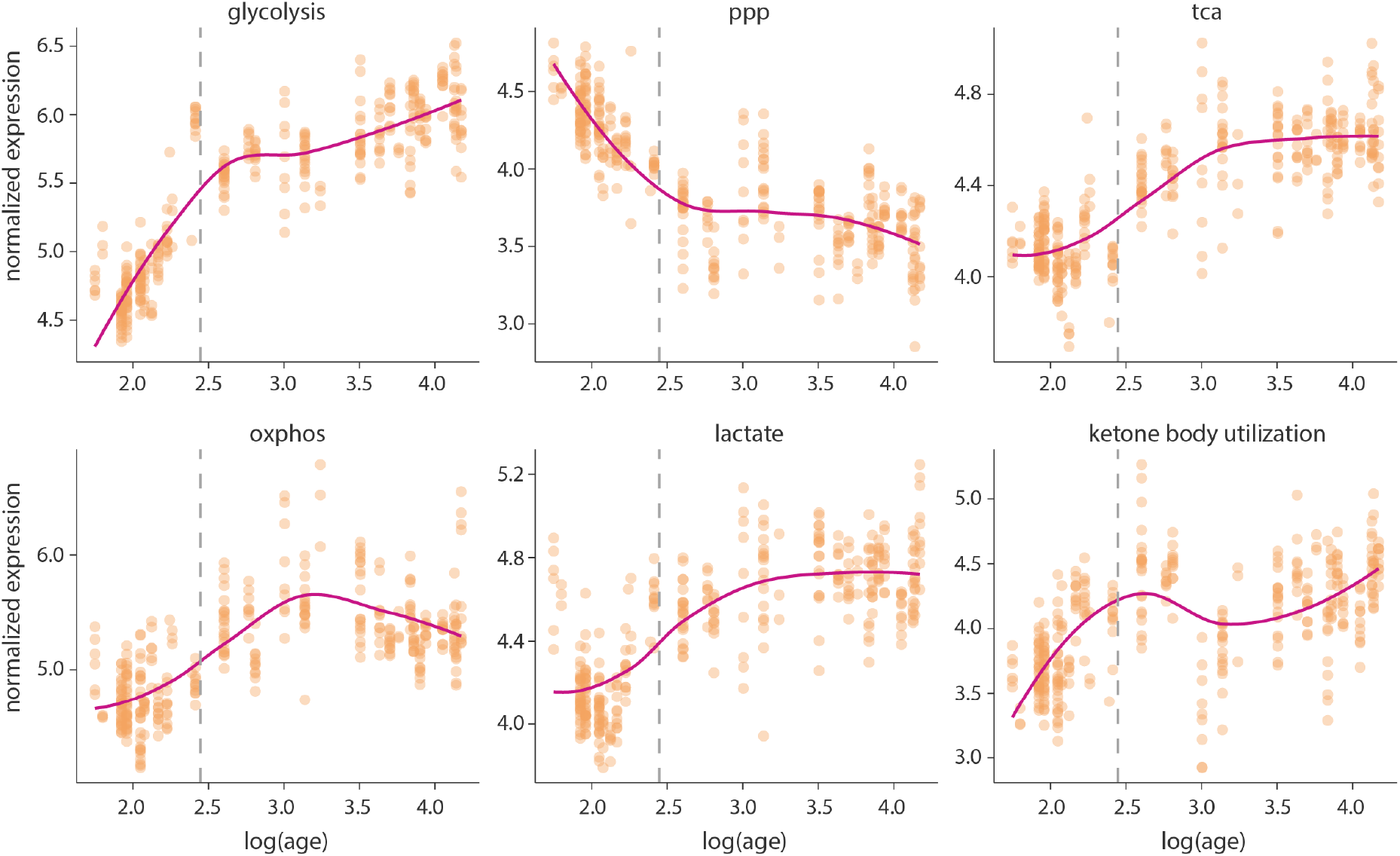
Lifespan trajectory of energy maps. Lifespan analysis was repeated using the non-parametric locally estimated scatterplot smoothing (LOESS) method. For each energy pathway, mean expression was calculated across all genes for each sample. The x-axis represent log_10_ transformed age in post conception days. Dots represent individual cortical samples at each age. The y-axis shows upper quartile normalized expression values. PPP, pentose phosphate pathway; TCA, tricarboxylic acid cycle; OXPHOS, oxidative phosphorylation.

**Figure S8.**
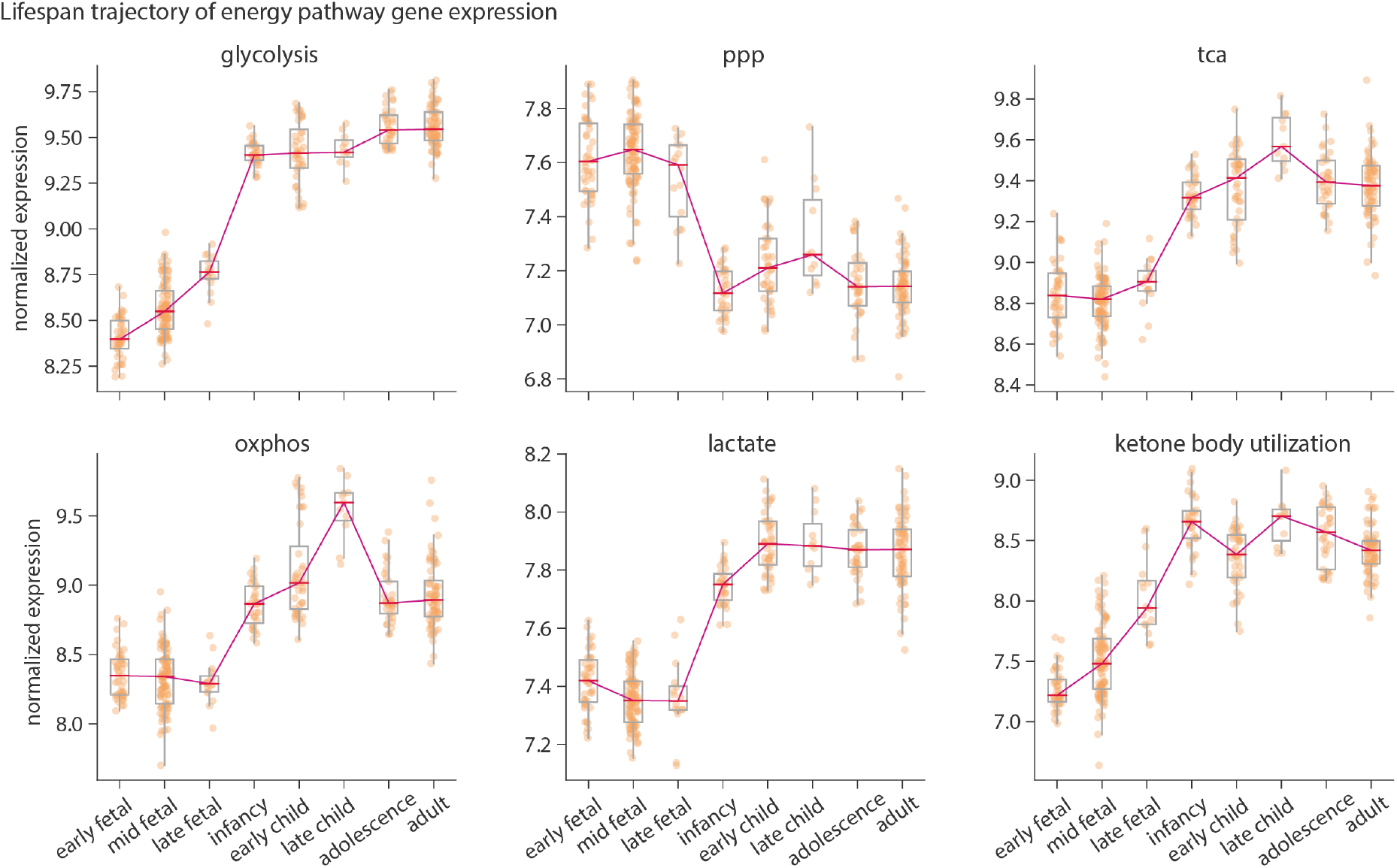
Microarray expression trajectory of energy maps across the lifespan. Lifespan analysis was repeated using the microarray data from the BrainSpan dataset [84]. For each energy pathway, mean expression was calculated across all genes for each sample. Samples were then grouped into eight age bins and the median expression across all samples falling into each age bin was calculated. Analysis only included cortical regions. The y-axis represents normalized log_2_(signal intensity) (see *Methods*). Dots represent individual samples in each age group. Line plot depicts the trajectory of median gene expression. For details of ages included in each group see Table S5. PPP, pentose phosphate pathway; TCA, tricarboxylic acid cycle; OXPHOS, oxidative phosphorylation.

**Figure S9.**
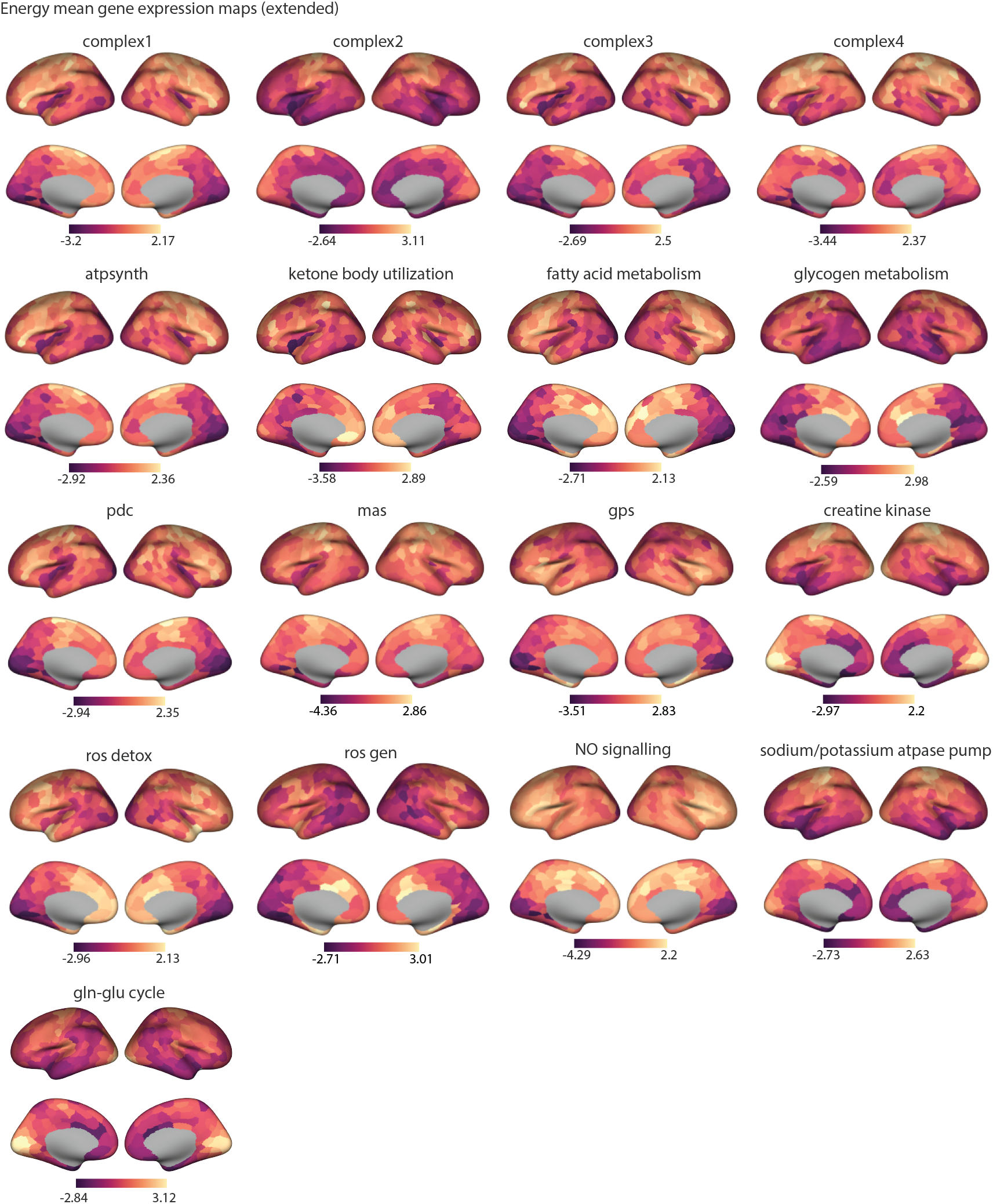
Extended set of energy relevant pathways. Mean gene expression maps were produced as before (see *Methods*). Color bars represent mean gene expression, z-scored across the 400 cortial regions in the Schaefer-400 parcelltaion. The clustering analysis of these maps can be found in Fig. S10. For completeness, the lifespan trajectory of the extended set of energy maps can be found in Fig. S12. atpsynth, ATP synthase complex; pdc, pyruvate dehydrogenase complex; mas, malate-aspartate shuttle; gps: glycerol-3-phosphate shuttle; ros detox, detoxification of reactive oxygen species; ros gen, generation of reactive oxygen species; no signaling, nitric oxide signaling; gln-glu cycle, glutamine-glutamate cycle.

**Figure S10.**
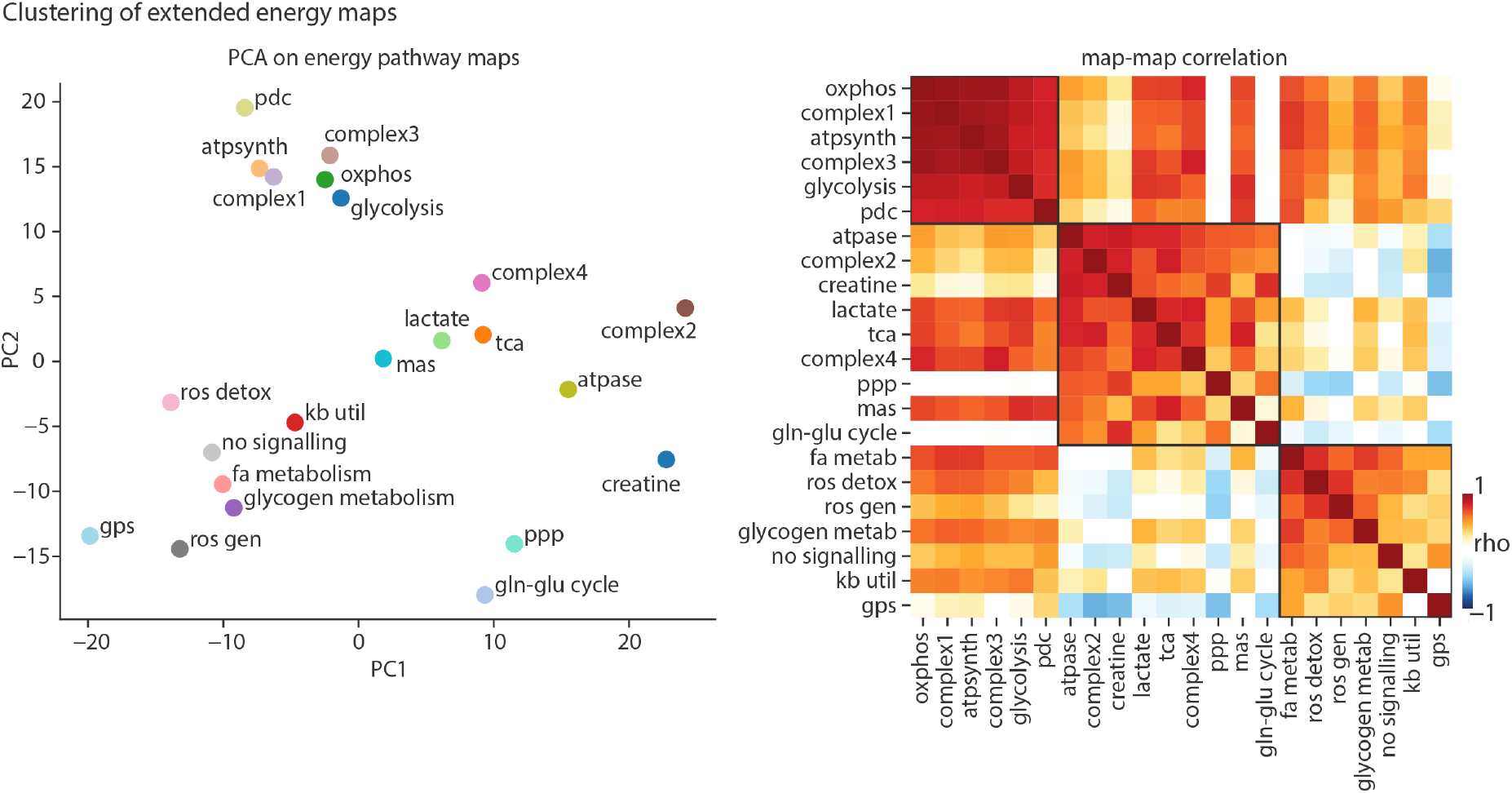
Clustering analysis of energy maps. The clustering among the complete set of energy maps was carried out using PCA (left), and Louvain modularity maximization algorithm (right). In the lefthand scatter plot, axes represent the first and second principal components. In the righthand heatmap, colors represent the pairwise Spearman’s correlations between energy maps. The first cluster groups together maps of glycolysis, OXPHOS, mitochondrial complex 1 and 3, the ATP synthase complex and the pyruvate dehydrogenase complex. This cluster therefore seems to capture the shared expression pattern between primarily energy-producing pathways responsible for glucose oxidation. The second cluster brings together the TCA, PPP, lactate, mitochondrial complexes 2 and 4, as well as creatine kinase gene expression and the malate-aspartate shuttle. The third cluster consists of fatty acid metabolism, ketone-body utilization and ROS related pathways. atpsynth, ATP synthase complex; pdc, pyruvate dehydrogenase complex; mas, malate-aspartate shuttle; gps: glycerol-3-phosphate shuttle; ros gen, generation of reactive oxygen species; no signaling, nitric oxide signaling; gln-glu cycle, glutamine-glutamate cycle.

**Figure S11.**
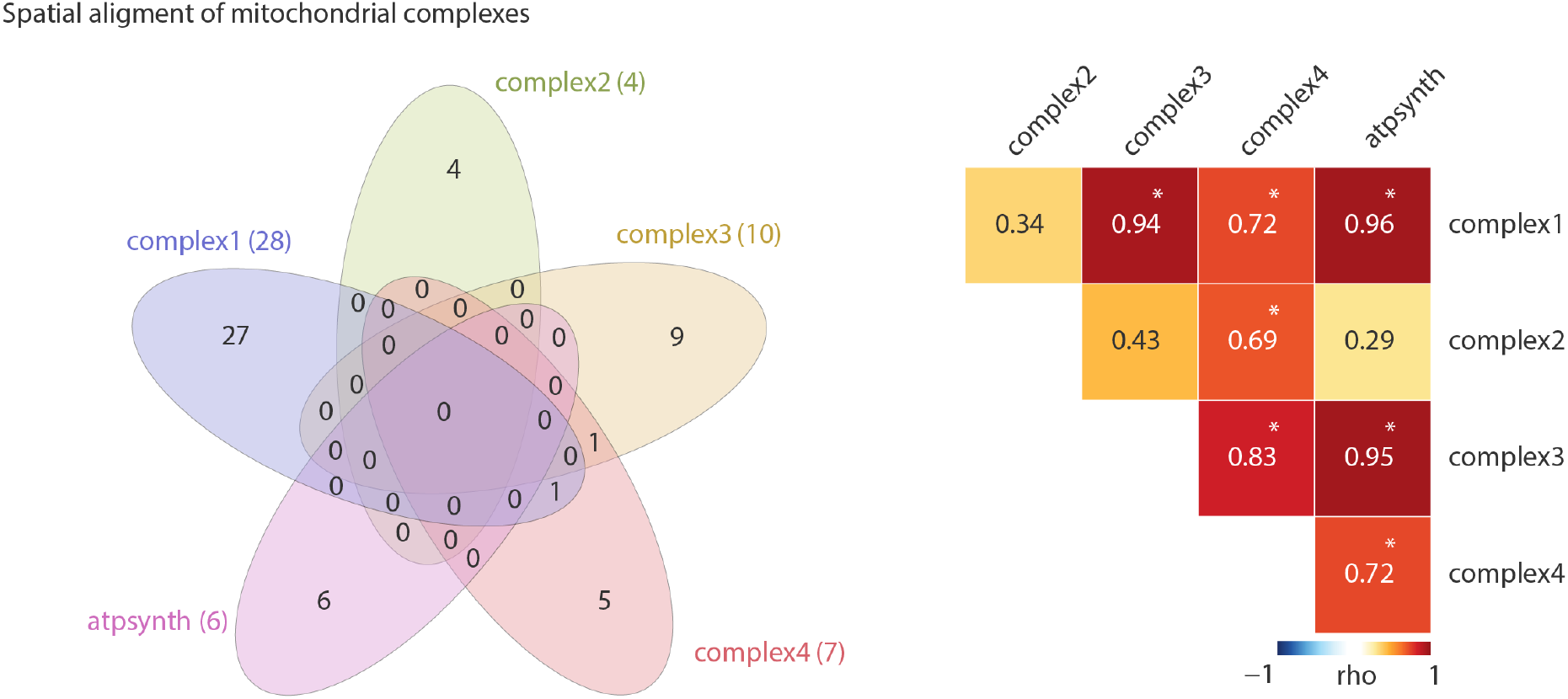
Alignment between the components of mitochondrial respiratory chain. Left: Venn diagram depicts the final number of genes in each mitochondrial complex and their overlap. Note that there is minimal overlap between genes in each complex. Right: Heatmap depicts pairwise Spearman’s correlations between mean gene expression maps. Gene sets for each mitochondrial complex were retrieved using GO and Reactome pathway IDs and mean expression maps where produced as before (see *Methods*). Asterisks show statistical significance when tested against a distribution of 10 000 correlations produced using spatial permutation testing. atpsynth, ATP synthase complex.

**Figure S12.**
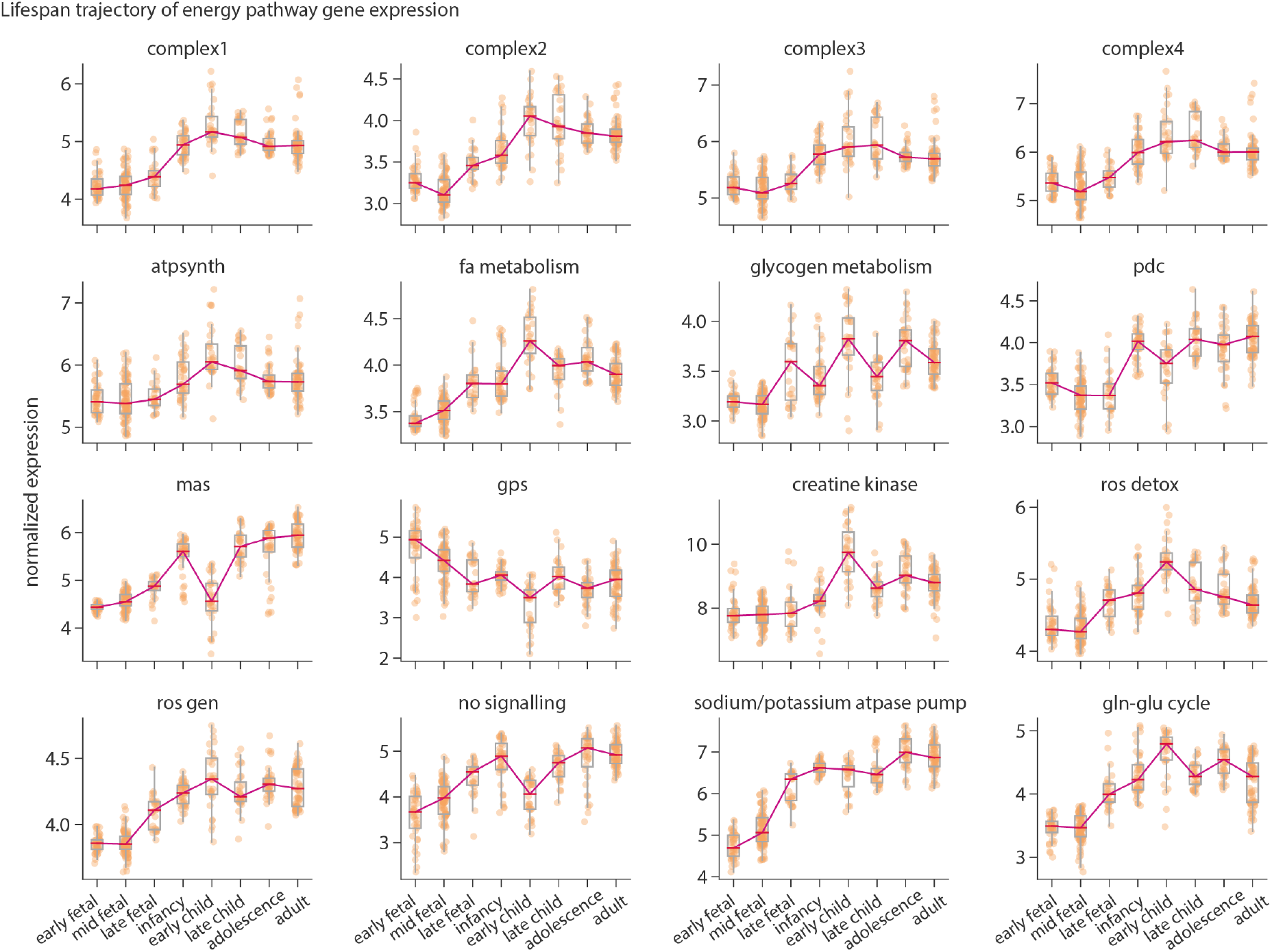
Lifespan trajectory of the extended energy metabolism maps. Developmental trajectories of energy metabolism pathways were produced using the BrainSpan RNA-seq expression dataset. For each energy pathway, mean expression was calculated across all genes for each sample and median expression across all samples falling into each age group was calculated. Analysis only included cortical regions. the y-axis represents normalized log_2_(RPKM) expression values. Dots represent individual samples in each age group. Line plot depicts the trajectory of median gene expression. atpsynth, ATP synthase complex; pdc, pyruvate dehydrogenase complex; mas, malate-aspartate shuttle; gps: glycerol-3-phosphate shuttle; ros detox: detoxification of reactive oxygen species; ros gen, generation of reactive oxygen species; no signalling, nitric oxide signaling; gln-glu cycle, glutamine-glutamate cycle.

**Figure S13.**
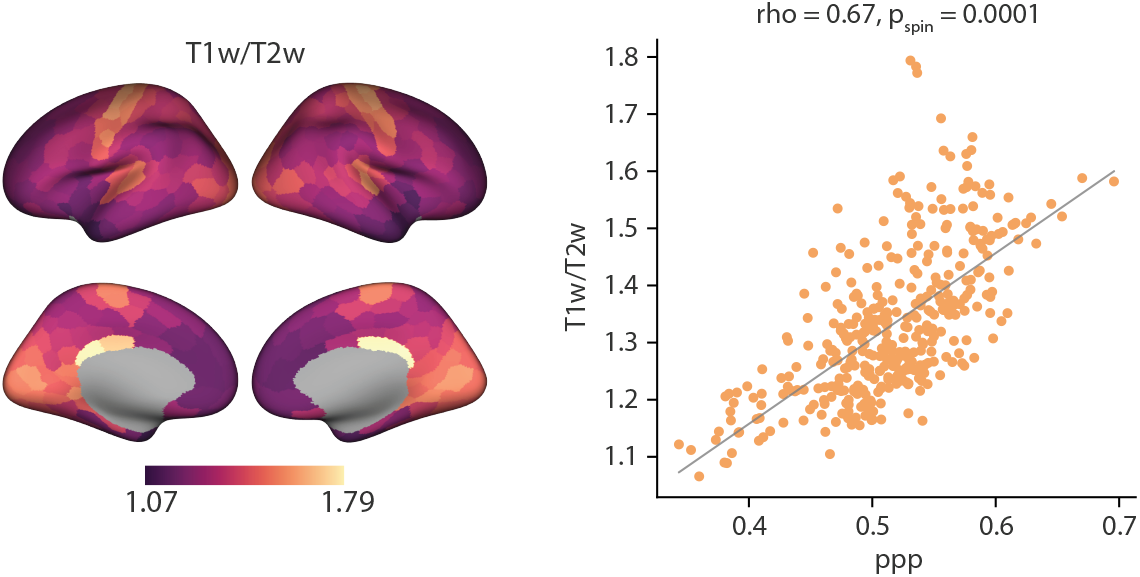
Alignment between the PPP and cortical T1w/T2w map. Spatial alignment between the maps were calculated using the Spearman’s correlation and tested against a distribution of 10 000 correlations produced from the spatial permutation nulls. The x-axis represents the PPP mean gene expression. Dots in each scatter plot represent the 400 cortical regions in the Schaefer-400 parcellation. ppp, pentose phosphate pathway; tca, tricarboxylic acid cycle.

**Figure S14.**
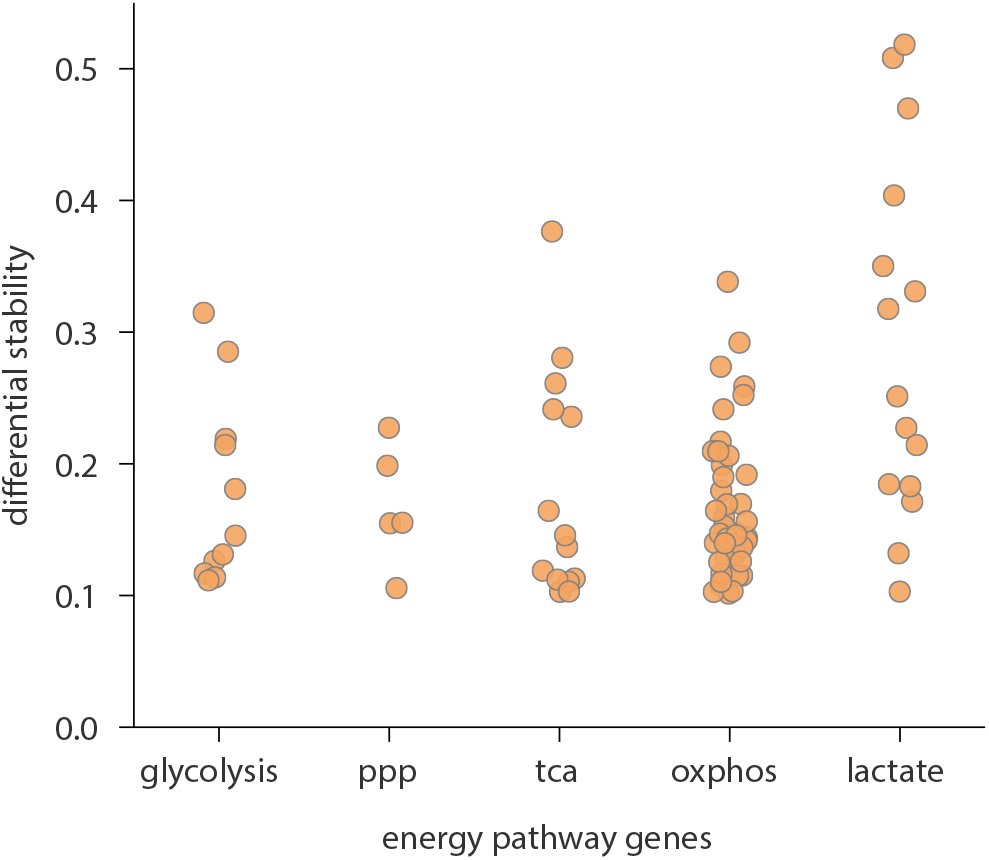
Differential stability of energy pathway genes. The differential stability distribution of the final energy pathway genes. Expression data were filtered to have a differential stability value >= 0.1. Dots represent individual genes in each pathway.

**TABLE S1.**
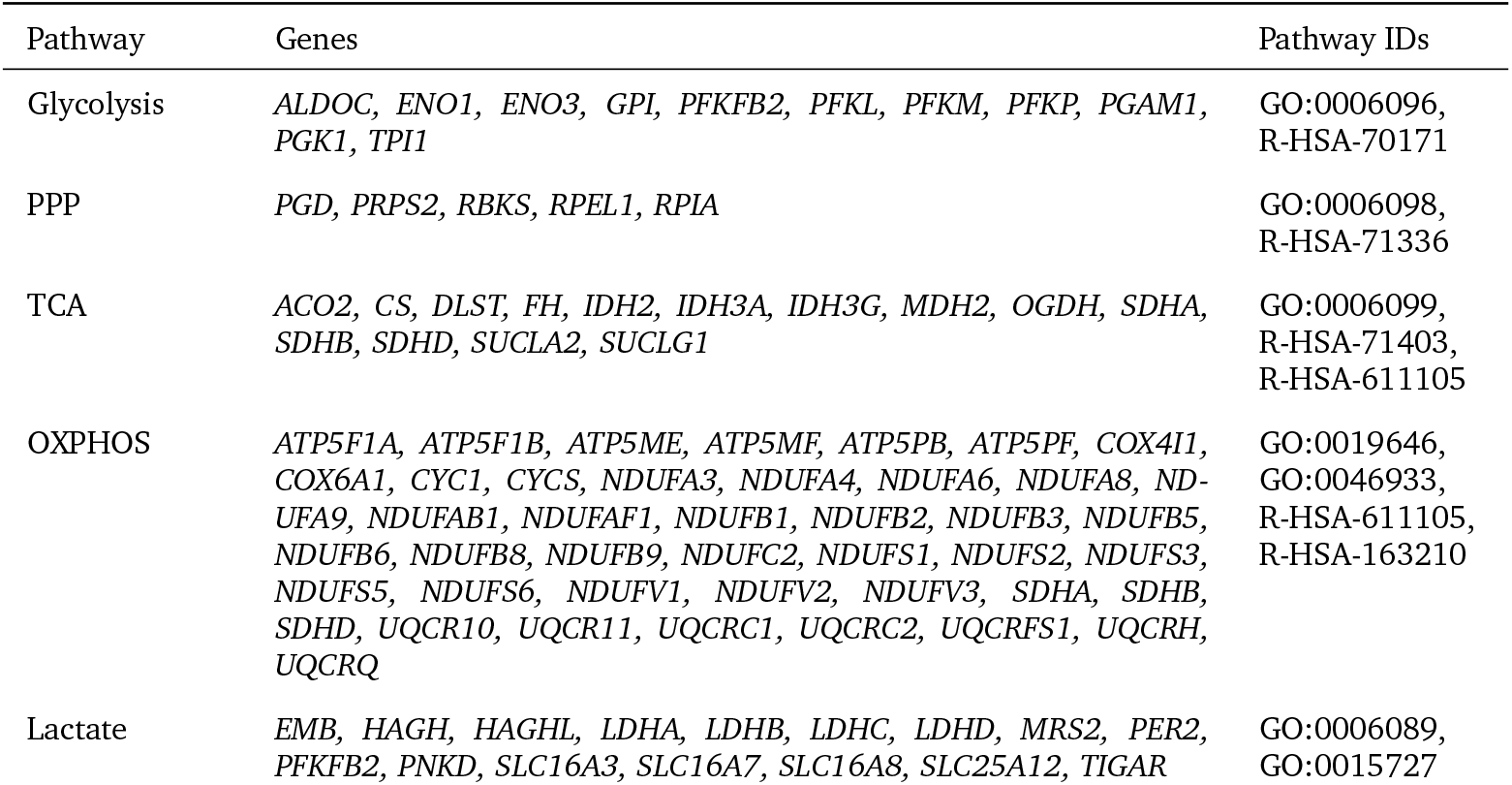
Energy metabolic pathway gene sets. Overview of energy metabolic pathways and their final gene sets included in this study. Gene sets were produced based on GO biological processes and Reactome pathway IDs. Genes annotated in both databases used for the analyses are listed. PPP, pentose phosphate pathway; TCA, tricarboxylic acid cycle; ETC, Electron transport chain; ATP, proton-coupled ATP synthesis.

**TABLE S2.** Visual ROIs. Visual regions in the Glasser atlas. Delineations were defined according to the supplementary neuroanatomical results from Glasser et al. [53] and https://neuroimaging-core-docs.readthedocs.io/en/latest/pages/atlases.html/.

| Label | Region abb. | Region name | Delineation |
| --- | --- | --- | --- |
| 1 | V1 | Primary Visual Cortex | Primary Visual |
| 2 | MST | Medial Superior Temporal Area | MT+ Complex and Neighboring Visual Areas |
| 3 | V6 | Sixth Visual Area | Dorsal Stream Visual |
| 4 | V2 | Second Visual Area | Early Visual |
| 5 | V3 | Third Visual Area | Early Visual |
| 6 | V4 | Fourth Visual Area | Early Visual |
| 7 | V8 | Eighth Visual Area | Ventral Stream Visual |
| 13 | V3A | Visual Area V3A | Dorsal Stream Visual |
| 16 | V7 | Seventh Visual Area | Dorsal Stream Visual |
| 17 | IPS1 | IntraParietal Sulcus Area 1 | Dorsal Stream Visual |
| 18 | FFC | Fusiform Face Complex | Ventral Stream Visual |
| 19 | V3B | Visual Area V3B | Dorsal Stream Visual |
| 20 | LO1 | Lateral Occipital Area 1 | MT+ Complex and Neighboring Visual Areas |
| 21 | LO2 | Lateral Occipital Area 2 | MT+ Complex and Neighboring Visual Areas |
| 22 | PIT | Posterior InferoTemporal complex | Ventral Stream Visual |
| 23 | MT | Middle Temporal Area | MT+ Complex and Neighboring Visual Areas |
| 138 | PH | Area PH | MT+ Complex and Neighboring Visual Areas |
| 152 | V6A | Visual Area V6A | Dorsal Stream Visual |
| 153 | VMV1 | VentroMedial Visual Area 1 | Ventral Stream Visual |
| 154 | VMV3 | VentroMedial Visual Area 3 | Ventral Stream Visual |
| 156 | V4t | Visual Area V4t | MT+ Complex and Neighboring Visual Areas |
| 157 | FST | Area FST | MT+ Complex and Neighboring Visual Areas |
| 158 | V3CD | Visual Area 3 c/d | MT+ Complex and Neighboring Visual Areas |
| 159 | LO3 | Lateral Occipital Area 3 | MT+ Complex and Neighboring Visual Areas |
| 160 | VMV2 | VentroMedial Visual Area 2 | Ventral Stream Visual |
| 163 | VVC | Ventral Visual Complex | Ventral Stream Visual |

**TABLE S3.** BrainSpan RNA-seq samples. Samples were binned into eight groups of major developmental stages [84]. Last column shows number of cortical samples used in the analysis. pcw, Post conception weeks; mos, months; yrs, years.

| Age group | Age | num. samples |
| --- | --- | --- |
| early fetal | 8 pcw, 9 pcw, 12 pcw | 40 |
| mid fetal | 13 pcw, 16 pcw, 17 pcw, 19 pcw, 21 pcw | 85 |
| late fetal | 24 pcw, 25 pcw, 26 pcw, 35 pcw, 37 pcw | 27 |
| infancy | 4 mos, 10 mos, 1 yrs | 41 |
| early childhood | 2 yrs, 3 yrs, 4 yrs | 30 |
| late childhood | 8 yrs, 11 yrs | 30 |
| adolescence | 13 yrs, 15 yrs, 18 yrs, 19 yrs | 36 |
| adulthood | 21 yrs, 23 yrs, 30 yrs, 36 yrs, 37 yrs, 40 yrs | 63 |

**TABLE S4.**
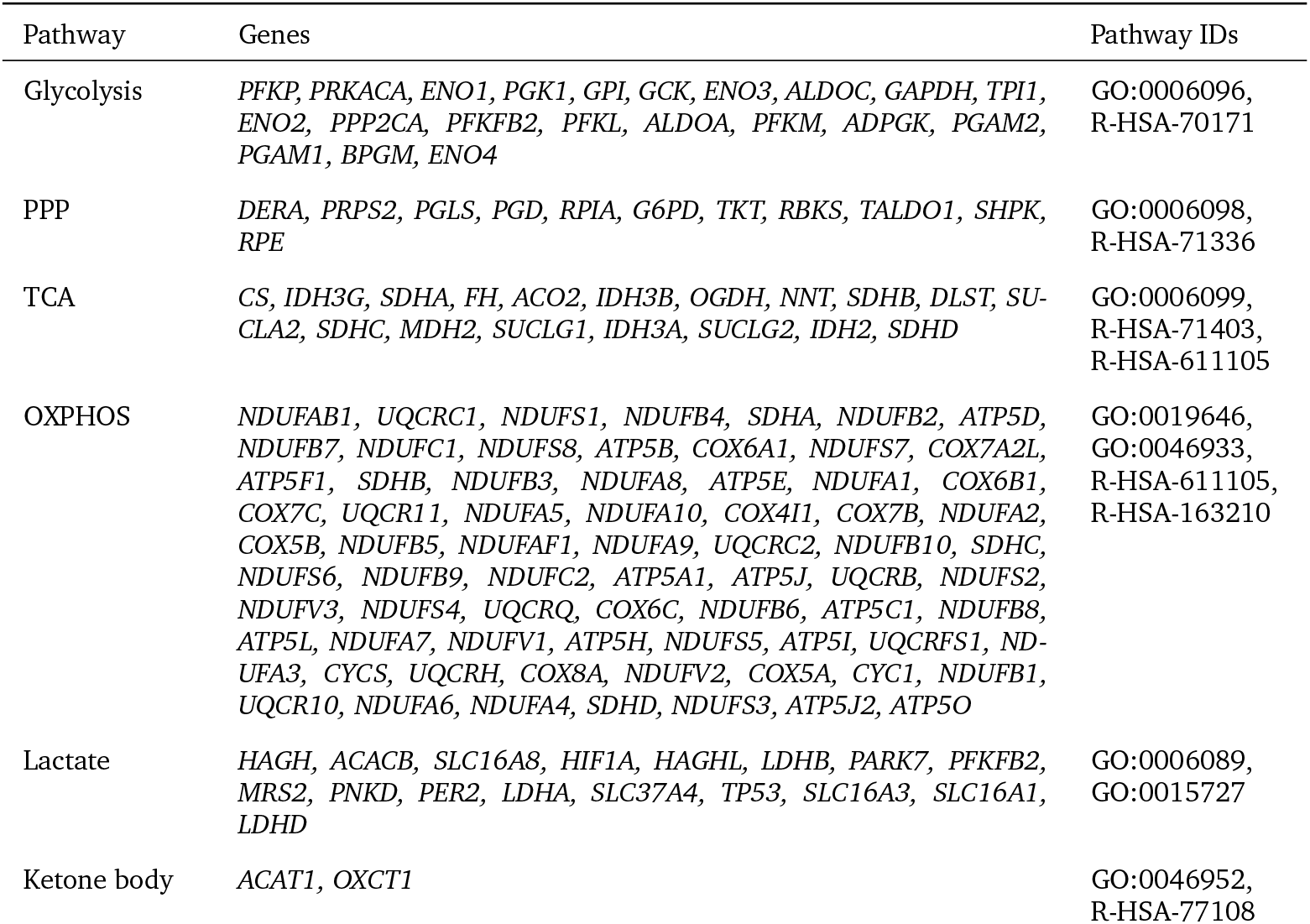
BrainSpan energy pathway gene sets. Energy metabolic pathways gene sets used in lifespan trajectory analysis. Note that the same GO biological processes and Reactome pathway IDs were used across all analysis and the difference in final gene sets for each pathway is the result of different gene data availability in AHBA and BrainSpan datasets. PPP, pentose phosphate pathway; TCA, tricarboxylic acid cycle; OXPHOS, oxidative phosphorylation; Lactate, lactate metabolism; Ketone Body, ketone body utilization.

**TABLE S5.** BrainSpan microarray samples. Samples were binned into eight groups of major developmental stages [84]. Last column shows number of cortical samples used in the analysis. pcw, Post conception weeks; mos, months; yrs, years.

| Age group | age | num.<br>samples |
| --- | --- | --- |
| early fetal | 8 pcw, 9 pcw, 12 pcw | 42 |
| mid fetal | 13 pcw, 16 pcw, 17 pcw, 19 pcw, 21 pcw | 95 |
| late fetal | 24 pcw, 25 pcw, 26 pcw | 16 |
| infancy | 4 mos, 10 mos, 1 yrs | 31 |
| early childhood | 2 yrs, 3 yrs, 4 yrs | 40 |
| late childhood | 8 yrs | 11 |
| adolescence | 13 yrs, 15 yrs, 18 yrs | 33 |
| adulthood | 21 yrs, 23 yrs, 30 yrs, 36 yrs, 37 yrs, 40 yrs | 66 |

